# Multiple pools of Protein Phosphatase 2A-B56 function to antagonize spindle assembly, promote kinetochore attachments and maintain cohesion in *Drosophila* Oocytes

**DOI:** 10.1101/2020.08.01.232512

**Authors:** Janet K. Jang, Amy C. Gladstein, Arunika Das, Zachary L. Sisco, Kim S. McKim

**Affiliations:** Waksman Institute, Rutgers, the State University of New Jersey, NJ-08854

## Abstract

Meiosis in female oocytes lack centrosomes, the major microtubule-organizing center, which makes them especially vulnerable to aneuploidy. In the acentrosomal oocytes of *Drosophila*, meiotic spindle assembly depends on the chromosomal passenger complex (CPC). Aurora B is the catalytic component of the CPC while the remaining subunits regulate its localization. Using an inhibitor of Aurora B activity, Binucleine 2, we found that continuous Aurora B activity is required to maintain the oocyte spindle during meiosis I, and this activity is antagonized by phosphatases acting on spindle associated proteins such as kinesins. Protein Phosphatase 2A (PP2A) exists in two varieties, B55 and B56. While both antagonize Aurora B, B55 has only minor roles in meiosis I spindle function. The B56 subunit is encoded by two partially redundant paralogs in the *Drosophila* genome, *wdb* and *wrd*. Knocking down both paralogs showed that the B56 subunit is critical for maintaining sister chromatid cohesion, establishing end-on microtubule attachments, and the metaphase I arrest in oocytes. We found that WDB recruitment to the centromeres depends on BUBR1, MEI-S332, and kinetochore protein SPC105R. While BUBR1 has been shown previously to stabilize microtubule attachments in *Drosophila* oocytes, only SPC105R is required for cohesion maintenance during meiosis I. We propose that SPC105R promotes cohesion maintenance by recruiting two proteins that recruit PP2A, MEI-S332, and the Soronin homolog Dalmatian.

## Introduction

Segregation errors during meiosis I or II results in aneuploidy in the zygotes that is often lethal. In humans, aneuploidy is a leading cause of infertility, spontaneous abortion, and birth defects, such as Down Syndrome, Klinefelter Syndrome, and Turner Syndrome (Kitajima, 2018; Mihajlovic and FitzHarris, 2018; Webster and Schuh, 2017). In females of many species, the meiotic spindle assembles in the absence of centrosomes and is directed largely by the chromosomes (Dumont and Desai, 2012; Radford et al., 2017). Prior studies have implicated two pathways in chromosome-directed spindle assembly: the chromosomal passenger complex (CPC) and the Ran pathway (Carazo-Salas et al., 2001; Carazo-Salas et al., 1999; Cesario and McKim, 2011; Colombié et al., 2008; Radford et al., 2012b). In *Drosophila*, the CPC is required for spindle and kinetochore assembly (Colombié et al., 2008; Radford et al., 2015; Radford et al., 2012b). Aurora B is the catalytic component while three other subunits, INCENP, Survivin and Borealin, control the localization of the CPC (Carmena et al., 2012; Trivedi and Stukenberg, 2020; van der Horst and Lens, 2014). Possible targets of the CPC include kinetochore proteins (Emanuele et al., 2008; Radford et al., 2015) and kinesin family proteins such as Subito and NCD, all of which contribute to assembly and organization of the meiotic spindle (Beaven et al., 2017; Jang et al., 2005). However, the relative contribution of positive and negative regulatory interactions in the formation of the meiotic acentrosomal spindle is poorly understood.

In many of its functions, Aurora B is antagonized by Protein Phosphatase I (PP1) or PP2A (Funabiki and Wynne, 2013; Saurin, 2018). The balance of kinase and phosphatase is important to regulate the steps of cell division such the initiation of prometaphase and the metaphase to anaphase transitions. Relative to mitotic cells, additional mechanisms regulating Aurora B – phosphatase antagonism are probably required for the unique properties of the two meiotic divisions (Keating et al., 2020). The first meiotic division involves the segregation of homologous chromosomes rather than sister chromatids. For the metaphase I – anaphase I transition, for example, sister centromeres must remain fused for them to co-orient and segregate to the same spindle pole (Wang et al., 2019; Watanabe, 2012). Furthermore, oocytes have arrest points and then are triggered by environmental signals to enter anaphase. For example, anaphase I in *Drosophila* females is triggered by changes in the oocyte environment that occur with passage through the oviduct (Heifetz et al., 2001; Horner and Wolfner, 2008). In mouse oocytes, PP2A-B56 regulates microtubule attachments based on timing of cell-cycle progression (Yoshida et al., 2015). Thus, the regulation of key meiotic events by phosphatases, such as spindle structure, sister chromatid cohesion and kinetochore-microtubule attachments in both fruit fly and mouse oocytes, may depend on developmental factors regulating progression through the meiotic divisions.

While Aurora B is required for initiating spindle assembly, it is not known if Aurora B activity is required to maintain the kinetochores and integrity of the spindle in oocyte meiosis. Using an inhibitor of Aurora B activity, Binucleine 2 (BN2) (Smurnyy et al., 2010), we observed the loss of all spindle microtubules in meiosis I oocytes. Thus, continuous Aurora B activity is required to maintain the spindle during meiosis I. These results also suggest that antagonism of Aurora B kinase by a phosphatase could regulate spindle dynamics. Indeed, we found that PP2A antagonizes Aurora B in the spindle maintenance function. PP2A is a is highly conserved heterotrimeric serine/threonine phosphatase composed of a scaffolding A subunit, a catalytic C subunit and one of multiple B subunits. The *Drosophila* genome like other organisms, encodes B subunits of two types, B55 and B56. Our results suggest both isoforms of PP2A antagonize the spindle assembly functions of Aurora B.

The two PP2A isoforms have different functions and target proteins. *Drosophila* PP2A-B55 has several targets many of which are phosphorylated by CDK1 and implicated in regulating the progression through G2 and mitosis (Kim et al., 2012; Rangone et al., 2011; Von Stetina et al., 2008; Wang et al., 2011b). In oocytes depleted of Twins, the *Drosophila* B55 subunit, meiotic entry and metaphase I proceeded surprisingly normal. In contrast, when the two *Drosophila* B56 isoforms, WRD and WDB, were depleted, there were dramatic defects in meiosis I. These results are consistent with results in *Drosophila* cell lines showing that loss of B56 has more severe defects in the events of metaphase and anaphase than loss of B55 (Chen et al., 2007).

We found that the PP2A-B56 WRD localizes to meiotic kinetochores and is required for sister chromatid cohesion and stabilizing attachments to microtubules. We also found that SPC105R, which we previously showed is required for sister centromere cohesion in meiosis (Radford et al., 2015; Wang et al., 2019), is required for PP2A-B56 localization. PP2A-B56 localization also depends on BubR1 and MEI-S332 (the *Drosophila* Shugoshin homolog), which is surprising because neither of these proteins are required for meiosis I cohesion. BUBR1/MEI-S332/PP2A-B56 maybe only one module that regulates meiotic cohesion. We propose that Dalmatian, the *Drosophila* orthologue of Soronin (Yamada et al., 2017), is part of a second module that has a cohesion protection function during meiosis I.

## Results

### Sustained Aurora B activity is required to maintain the oocyte meiotic spindle

In the *Drosophila* ovary, the nuclear envelope breaks down and spindle assembly begins in stage 13 oocytes (Gilliland et al., 2009). By stage 14, a bipolar spindle forms. In previous work we found that the chromosomal passenger complex (CPC), through the activity of Aurora B kinase, is required for assembly of a bipolar spindle in *Drosophila* oocytes (Radford et al., 2012b). In these experiments, germline-specific RNAi was used to deplete Aurora B or the targeting subunit INCENP prior to nuclear envelop breakdown. To determine the effect of depleting Aurora B activity during prometaphase, we treated stage 14 oocytes with the drug Binucleine 2 (BN2), which inhibits the kinase activity of Aurora B (Eggert et al., 2004; Smurnyy et al., 2010). Spindle assembly was monitored with immunofluorescence for tubulin, kinetochore protein SPC105R/KNL1, central spindle protein Subito/MKLP2 and CPC proteins INCENP or Aurora B. In control experiments, when wild-type females were treated with DMSO only, 100% of oocytes assembled a bipolar spindle that included Subito and the CPC within the central anti-parallel microtubules (Figure 1A, B). In wild-type oocytes treated with 50 µM BN2 for one hour, the spindle was absent or faint in 85% of oocytes. Only 15% of oocytes had robust bipolar spindles (Figure 1A). These results show that maintaining the meiotic spindle depends on Aurora B activity.

**Figure 1:**
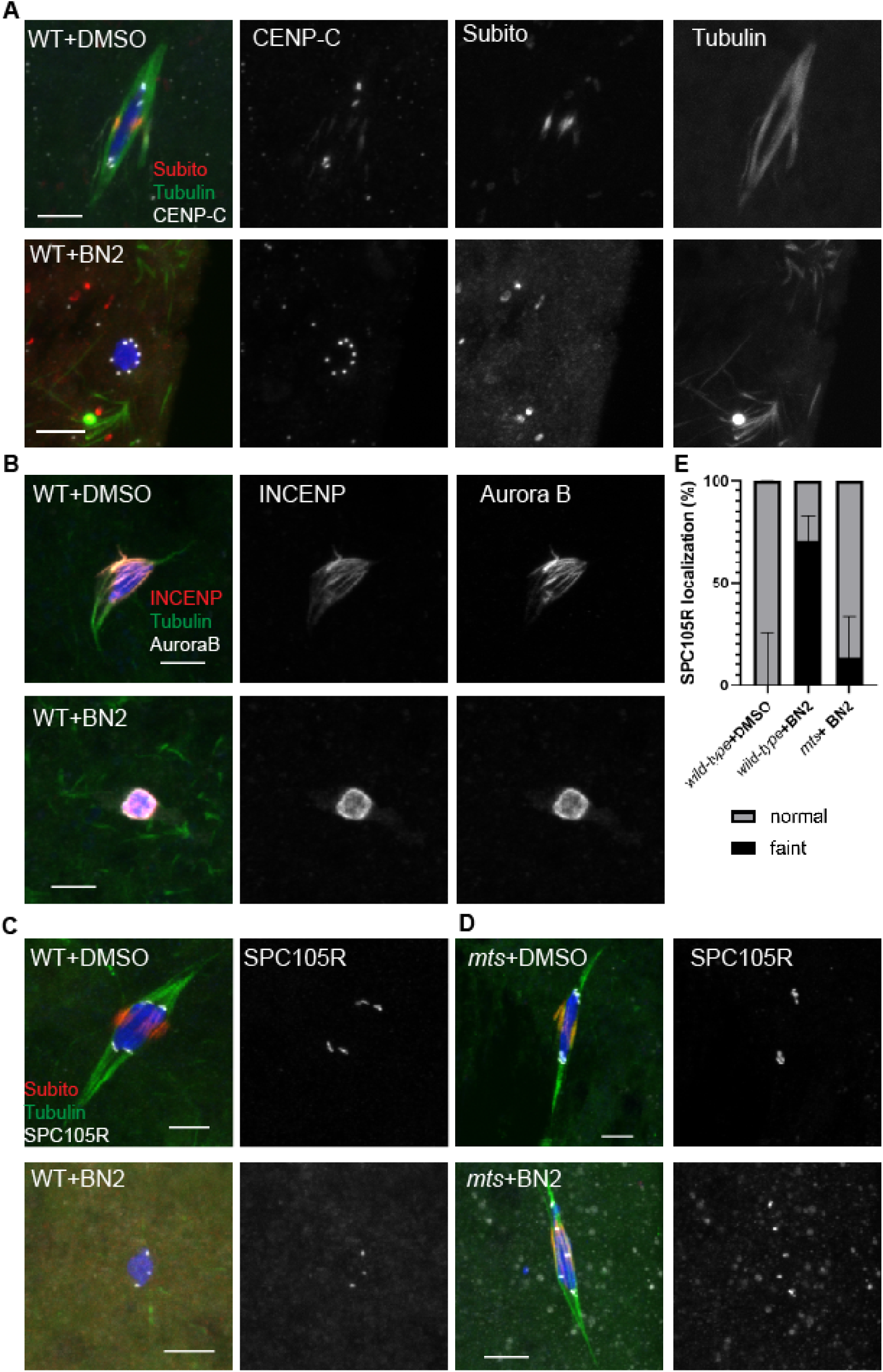
Constant Aurora B activity is required for maintaining spindle microtubules and SPC105R localization in oocytes. (A) Wild-type (WT) oocytes treated with either 0.001% DMSO or 50µM BN2. DNA is shown in blue, Subito in red and tubulin in green. All images are maximum projections of Z-stack and scale bars are 5 µm. (B) Wild-type oocytes treated with either 0.001% DMSO or 50µM BN2 immunostained with DNA (blue), INCENP (red), tubulin (green) and Aurora B (white). (C) Localization of kinetochore protein SPC105R (white) was reduced in wild-type oocytes treated with BN2 (72% faint and reduced localization, n=36) compared to controls (0% faint, n=11). (D) SPC105R localization was retained in *mts*^*HMJ22483*^ RNAi oocytes treated with BN2 (16% weak, n=19, P<0.0001, Fisher’s exact test). E) Summary of SCP105R localization data.

In the absence of Aurora B, INCENP localizes to the meiotic chromosomes (Radford et al., 2012b). BN2 treatment allowed us to test if kinase activity is required for Aurora B localization. When Aurora B activity was inhibited with BN2, INCENP localized to the chromosomes, showing that upon BN2 treatment, the CPC moves from the microtubules to the chromosomes (Figure 1B). In addition, Aurora B localized to the chromosomes, demonstrating that inhibition of phosphorylation by BN2 did not prevent Aurora B association with INCENP and localization to the chromosomes. To directly test Aurora B activity, we used an antibody for one of its targets, phosphorylation of INCENP (pINCENP) at a conserved serine in the C-terminal domain (Salimian et al., 2011; Wang et al., 2011a). While solvent treated controls had robust pINCENP in the central spindle, the chromosome localized INCENP in BN2 treated oocytes had lost its phosphorylation (Figure 2A,B). These results show that Aurora B phosphorylation of INCENP is not required for chromosome localization of the CPC.

**Figure 2:**
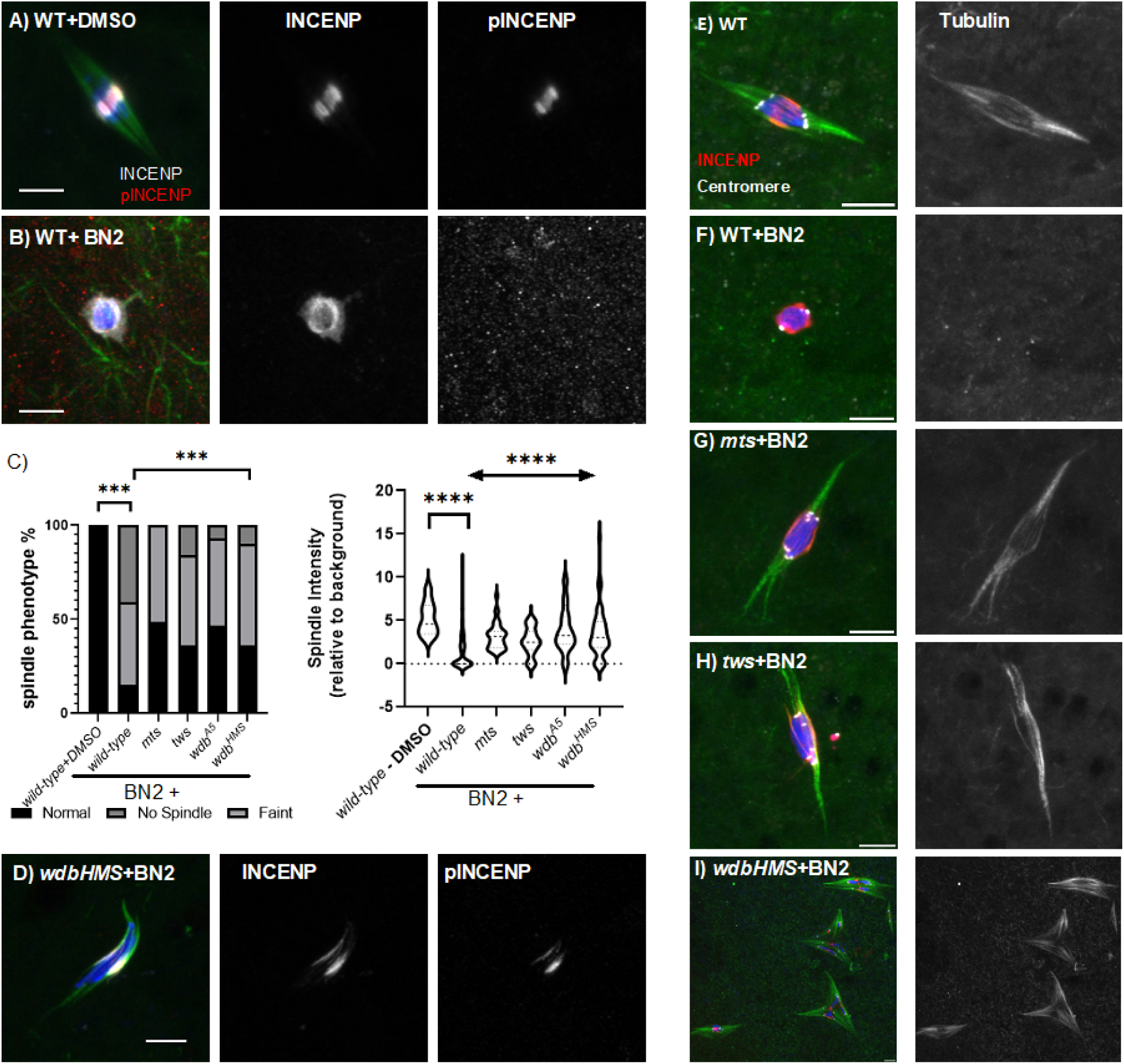
PP2A antagonizes Aurora B activity. Oocytes treated with (A) 0.001% DMSO had pINCENP (n=19/20) while (B) oocytes treated with 50µM BN2 had pINCENP at a reduced frequency (6/15, p=0.007). The BN2 treated oocytes with pINCENP tended to have some residual spindle assembly. Phosphorylated INCENP (pINCENP) is shown in red, INCENP in white, tubulin in green and DNA in blue. (C) Qualitative and quantitative assessment of spindle assembly in wild-type+DMSO (n=11), wild-type + BN2 (n=84), *mts* (n=39), *tws* (n=25), *wdb*^*A5*^ (n=30) and *wdb*^*HMS01864*^ (n=50) RNAi oocytes treated with BN2. (D) Retention of pINCENP when *wdb* RNAi oocytes were treated with 50µM BN2 (n=15). (E -I) Wild-type and RNAi oocytes treated with 50µM BN2, showing INCENP (red), centromeres (white), tubulin (green), and DNA (blue). Tubulin is shown in the right panel. Scale bars are 5μm.

The *Drosophila* oocyte meiotic spindle has two types of microtubules based on the nature of their + ends. The kinetochore microtubules have their + ends at a kinetochore. The anti-parallel microtubules have their + ends within the central spindle (Jang et al., 2005; Radford et al., 2015). Subito, a Kinesin-6 required for interpolar microtubule assembly of the central spindle (Jang et al., 2005), was lost upon BN2 treatment (Figure 1A), demonstrating Subito localization depends on Aurora B activity and/or microtubules. SPC105R, which is required for kinetochore microtubules (Radford et al., 2015), was also reduced, although not eliminated (Figure 1C,E). These results show that maintenance of the two major organizers of oocyte spindle microtubules depends on sustained Aurora B activity.

### PP2A antagonizes the spindle assembly function of the CPC

BN2 treatment was applied during prometaphase, and therefore, after the initial phosphorylation of CPC substrates and assembly of spindles. To explain why inhibiting CPC activity causes loss of spindle microtubules, we tested the hypothesis that phosphatases (PP1 or PP2A) dephosphorylate Aurora B targets. To identify the phosphatase responsible for antagonizing Aurora B, shRNA expression under the control of the UAS/Gal4 system was used for germline specific RNAi of each phosphatase. Each shRNA was expressed using *mata4-GAL-VP16*, which induces expression UAS controlled transgenes after pre-meiotic DNA replication but throughout most stages of meiotic prophase during oocyte development (Radford et al., 2012b). During this time, the oocyte grows and matures, but does not divide, while the target gene expression is depleted. We refer to oocytes expressing an shRNA using *mata4-GAL-VP16* as “RNAi oocytes”.

PP1-87B is the major PP1 isoform functioning in oocytes (Wang et al., 2019). *Pp1-87B* RNAi oocytes were treated with BN2 and spindle disintegration was measured. If PP1-87B opposes the Aurora B spindle assembly function in oocytes, then addition of BN2 after depletion of PP1-87B would not cause loss of spindle microtubules. However, *Pp1-87B* RNAi oocytes treated with BN2 had reduced spindle microtubules in a significant number of oocytes when compared to the *Pp1-87B* RNAi solvent treated controls (Figure S 1). SPC105R localization and some kinetochore microtubule fibers were retained in *Pp1-87B* RNAi oocytes treated with BN2, however, the central spindle and Subito localization was always absent (Figure S 1). These observations are consistent with our previous observations that *Pp1-87B* RNAi oocytes are partially resistant to BN2 treatment because loss of PP1-87B stabilizes kinetochore-microtubule (KT-MT) attachments (Wang et al., 2019).

To test if PP2A has a role in spindle assembly, we depleted specific subunits by RNAi. PP2A exists in multiple complexes, although all the complexes contain the same A (PP2A-29B) and catalytic C (MTS = Microtubule star) subunits. Depletion of the A-subunit (*Pp2A-29B, GLC01651*) or C-subunit (*mts, HMS04478*) of PP2A resulted in no mature stage 14 oocytes, indicating that PP2A activity is required for oocyte development. However, expression of one shRNA for *mts* (*HMJ22483*) produced stage 14 oocytes, probably due to partial depletion of MTS protein in *HMJ22483* RNAi oocytes (Table 1). When *mts*^*HMJ22483*^ RNAi oocytes were treated with BN2, a bipolar spindle and SPC105R localization was retained in most oocytes (Figure 1D, E, Figure 2C,E-G). Unlike *Pp1-87B* RNAi oocytes, Subito localization was retained (Figure 1C). These results suggest that PP2A antagonizes the role of the CPC in maintaining the kinetochores and the microtubules of the bipolar spindle.

**Table 1:**
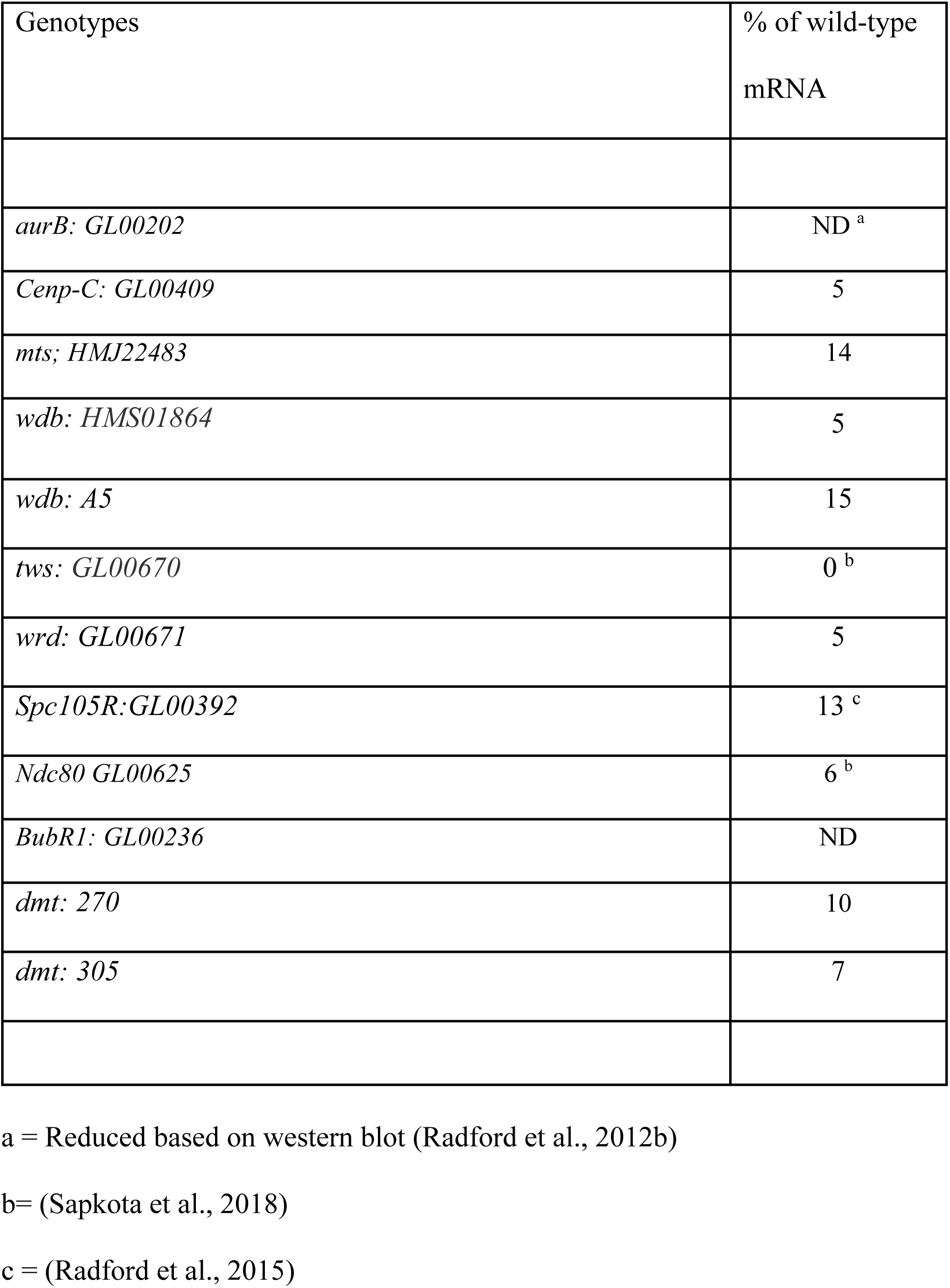
Summary of RNAi lines.

### Both PP2A complexes antagonize the role of the CPC in spindle maintenance

PP2A exists in at least two complexes, and in *Drosophila* contains either a B55 (Twins=TWS) or one of two B56 (Widerborst=WDB or Well-rounded = WRD) subunits (Chen et al., 2007). In order to determine which PP2A complex opposes the CPC, BN2 treatment was done in the presence of B subunit RNAi. A shRNA line for *tws* (*GL00670*) caused complete sterility when expressed using *mataGal4* and reduced the mRNA to 0% of wild-type levels (Table 1). In *tws* RNAi oocytes treated with BN2, oocyte spindle assembly was restored in ∼80% of oocytes (Figure 2C, H). For the B56 subunit, we focused on WDB because it is essential and WRD is not (see below). Two shRNA lines were used for *wdb*: one generated by TRiP (Ni et al., 2011) (*HMS01864*) and one generated in our laboratory (*A5*). Both shRNAs efficiently knocked down *wdb* RNA (Table 1) and the oocytes had similar phenotypes. In both *wdb*^*HMS01864*^ and *wdb*^*A5*^ RNAi oocytes treated with BN2, the spindle was present in ∼80% of oocytes (Figure 2C, I). Depletion of WDB also restored localization of INCENP and phospho-INCENP to the central spindle (Figure 2D). These results demonstrate that both B55 and B56 PP2A complexes antagonize Aurora B activity in the process of spindle assembly and maintenance.

### Kinesin 13 Klp10A is a CPC and PP2A target

The maintenance of the meiotic spindle could depend on Aurora B inhibiting depolymerizing factors or promoting spindle assembly factors. Thus, PP2A could promote the removal of tubulin subunits from the spindle, while Aurora B could promote addition of tubulin subunits to the spindle. Therefore, we used FRAP to test the hypothesis that PP2A promotes spindle dynamics. However, in *mts* RNAi oocytes, we found tubulin turnover to be unchanged from wild-type (Figure S 2). Thus, PP2A is not required for the turnover of microtubule subunits within the spindle.

The kinesin-13 homolog KLP10A depolymerizes spindle microtubules in oocytes (Radford et al., 2012a). To test the hypothesis that PP2A regulates spindle depolymerizing factors, we examined meiotic spindles in *Klp10A* RNAi oocytes treated with BN2. In *Klp10A* RNAi control oocytes, the spindle is very long and disorganized (Figure 3B). In contrast, the spindle was retained in BN2 treated *Klp10A* RNAi oocytes (Figure 3C), suggesting that KLP10A is a possible CPC and PP2A target and that the maintenance of the spindle could depend on inhibiting depolymerizing factors. The spindle phenotype of BN2 treated *Klp10A* RNAi oocytes was more variable than BN2 treated wild-type or PP2A RNAi oocytes. In some oocytes, central spindle and kinetochore microtubules appeared to be intact. In other oocytes, the spindle was retained but Subito or INCENP localization was abnormal, indicating that the central spindle and/or kinetochore microtubules were absent (Figure 3D). These results suggest KLP10A is inactivated by CPC phosphorylation, and activated when dephosphorylated by PP2A, although the central spindle and kinetochores maybe more sensitive to loss of Aurora B kinase activity than microtubule bundles in general.

**Figure 3:**
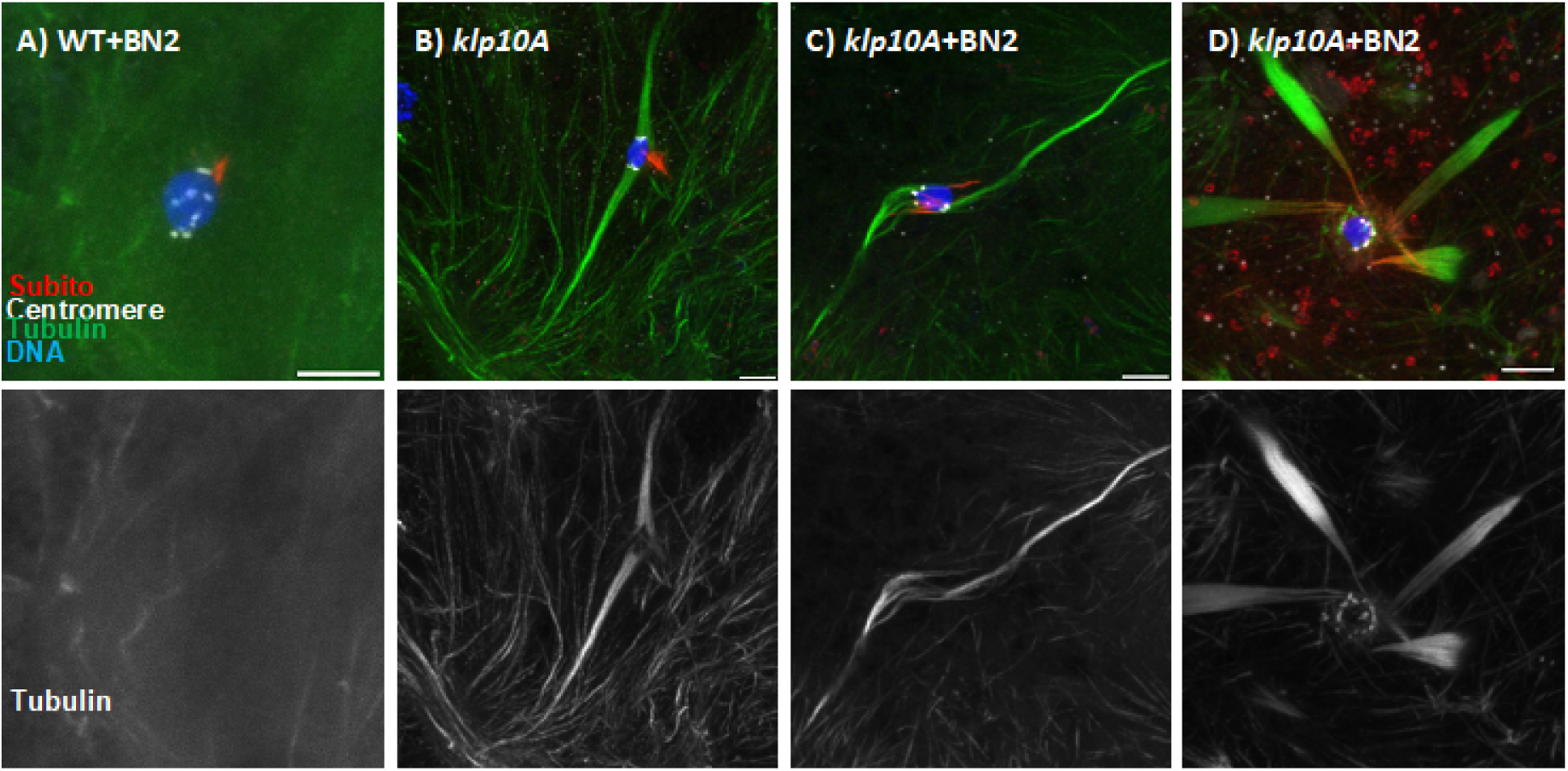
Aurora B antagonizes KLP10A. (A) Wild-type (WT) oocytes treated with BN2. (B) Untreated *klp10A* RNAi oocyte showing characteristic long spindle phenotype. (C) Most wild-type oocytes treated with BN2 failed to form a spindle (16/20). Most *Klp10A RNAi* oocytes treated with BN2 developed a long spindle (39 out of 45, p<0.0001). (D) In some cases, the spindle in *Klp10A RNAi* oocytes treated with BN2 was detached from the chromosomes, as if the kinetochore attachments were destabilized. The Images are shown with centromeres in white, Tubulin in green, Subito in red, and DNA in blue. Tubulin channel shown in bottom panel. Scale bars are 5μm.

### WRD compensates for loss of WDB in the germline

*Drosophila* has two B56-type paralogs, WDB and WRD. WDB and WRD share 68% identity with long stretches of identity. Null mutants of *wrd* are viable and fertile (Hahn et al., 2010; Moazzen et al., 2009). An shRNA targeting *wrd* (*GL00671*) substantially reduced mRNA levels (Table 1). Consistent with the phenotype of *wrd* mutants, females expressing *GL00671* in the germline were fertile and nondisjunction was rare (Table 2). In contrast, ubiquitous expression of *wdb* shRNA caused lethality (Methods), similar to homozygous null mutations of *wdb*. Thus, WDB appears to be the more important B56 subunit. However, the germline *wdb* RNAi phenotypes were milder than expected. Maternal expression of either *wdb* shRNA caused reduced fertility but the females were not sterile and meiotic nondisjunction was low (Table 2). The spindle structure of *wdb* RNAi oocytes was similar to wild-type; they were bipolar, and assembled a robust central spindle (Figure 4A). Furthermore, bundles of microtubules terminated at the kinetochores, and the centromeres were oriented towards a pole, suggesting MT-KT attachments were forming properly. Similar observations were made with *tws* RNAi oocytes. These data suggest that TWS and WDB are not essential for bipolar spindle assembly.

**Table 2:** Fertility of RNAi females

| Genotype | Number of Progeny | Number of Vials | Average Progeny per Vial |
| --- | --- | --- | --- |
| *mts<sup>HMJ22483</sup>/mata* | 0 <sup>a</sup> | 20 | 0 |
| *mts<sup>HMS04478</sup>/mata* | 0 <sup>b</sup> | 10 | 0 |
| *wdb<sup>HMS01864</sup>/mata* | 257 | 10 | 25.7 |
| *wrd<sup>GL00671</sup>/mata* | 1440 | 20 | 72 |
| *wdb<sup>A5</sup>/mata* | 216 | 33 | 7 |
| *wdb<sup>HMS01864</sup>/nos* | 1044 | 10 | 104 |
| *wdb<sup>A5</sup>/nos* | 533 | 10 | 53 |
| *wrd<sup>-</sup>; wdb<sup>HMS01864</sup>/mata* | 4 | 14 | 0.3 |
| *wrd<sup>GL00671</sup>; wdb<sup>HMS01864</sup>/mata* | 0 <sup>a</sup> | 15 | 0 |
| *wrd<sup>GL00671</sup>; wdb<sup>HMS01864</sup>/nos* | 0 <sup>b</sup> | 15 | 0 |
| *dmt<sup>270</sup>/nos* | 381 | 9 | 42 |
| *dmt<sup>305</sup>/nos* | 0 <sup>a</sup> | 30 | 0 |
| *dmt<sup>270</sup>/mata* | 0 <sup>a</sup> | 20 | 0 |
| *dmt<sup>305</sup>/mata* | 0 <sup>a</sup> | 20 | 0 |

a: Stage 14 oocytes present
b: No Stage 14 oocytes

**Figure 4:**
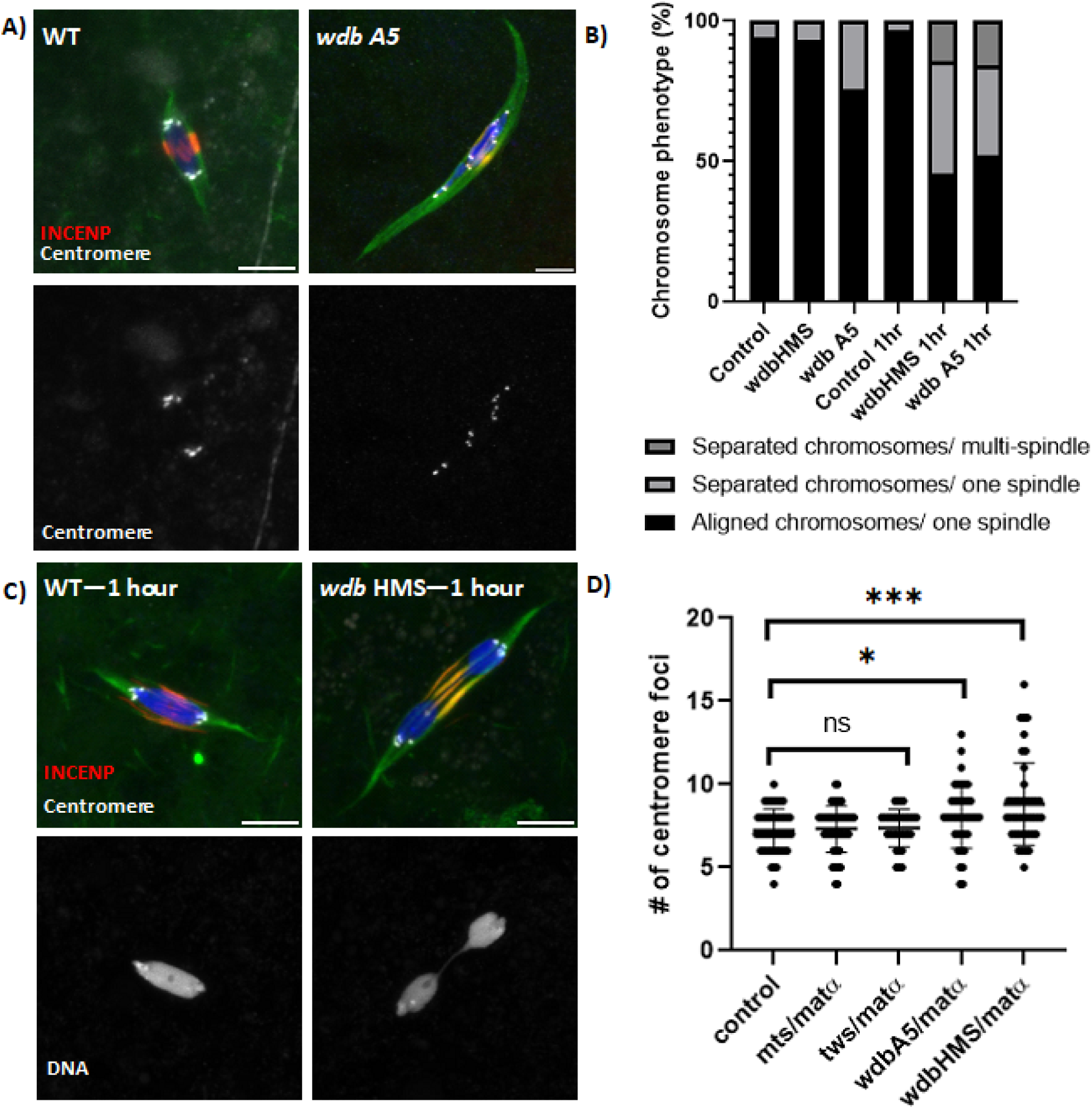
Spindle assembly in PP2A RNAi oocytes. (A) Wild-type (WT) or wd*b*^*A5*^ RNAi oocytes with INCENP in red, centromeres in white, tubulin in green and DNA in blue. The bottom panel shows the centromeres. The scale bars are 5μm. (B) Percent chromosome phenotype for each genotype (control n=51; *wdb*^*HMS01864*^ n=30; *wdb*^*A5*^ n=41; control *1 hr* n=32. *wdb*^*A5*^ 1 hr n=35; *wdb*^*HMS01864*^ 1 hr n=25). Precocious anaphase is indicated by the separation of chromosomes towards the poles. (C) Wild-type or *wdb*^*HMS01864*^ oocytes incubated in Robbs buffer for 1 hour without BN2. Bottom panel shows the DNA channel. (D) Plot showing the number of centromere foci for each genotype (control n=51; *tws* n=32; *mts* n=50; *wdb*^*A5*^ n=39; *wdb*^*HMS01864*^ n=43).

WDB and WRD are partially redundant in mitosis (Chen et al., 2007). To test for redundancy, two genotypes were generated to reduce expression of both WDB and WRD: females expressing shRNA to both *wdb* and *wrd* (*wrd*^*GL00671*^; *wdb*^*HMS01864*^), and females expressing *wdb* shRNA and hemizygous for a *wrd* mutation (Materials and Methods) (*wrd*^*-*^ ; *wdb*^*HMS01864*^). When using *mata4-GAL-VP16* for expression of the shRNA, the double knockdown females were completely sterile, unlike the single RNAi females. In addition, when using *nos-GAL4-VP16*, which promotes expression of the shRNA in premeiotic germ cells, no oocytes were produced, also unlike the single RNAi females. These results demonstrate that WRD and WDB are partially redundant in the mitotic and meiotic germline and expression of WRD is sufficient for fertility. In contrast, expression of WDB, but not WRD, is sufficient for viability.

### PP2A-B56 protects meiotic sister chromatid cohesion and promotes end-on microtubule attachments

*Drosophila* oocytes arrest in metaphase I at developmental stage 14, and do not proceed into anaphase I and meiosis II until activated by passage through the oviduct (Heifetz et al., 2001). Stage 14 oocytes can also be induced to proceed past metaphase I by incubation in certain buffers (Endow and Komma, 1997; Page and Orr-Weaver, 1997). Our methods use a modified Robbs’ buffer to prevent premature activation (Theurkauf and Hawley, 1992). Indeed, most wild-type and *wdb* RNAi oocytes were arrested in metaphase I (Figure 4A, B). Furthermore, even after control oocytes were incubated for one hour in modified Robbs, metaphase I arrest was usually maintained (Figure 4B, C). We were surprised, therefore, to find that the one-hour incubation in modified Robbs buffer with BN2 treatment induced precocious anaphase in *wdb* RNAi oocytes (Figure 2I). To investigate if this was an effect of the BN2 treatment, *wdb* RNAi oocytes were incubated in modified Robbs buffer for one hour without BN2. In these conditions, precocious separation of homologous chromosomes was observed in approximately 50% of oocytes (Figure 4B, C). These phenotypes were present in *wdb* RNAi oocytes but not in *tws* RNAi oocytes. These data suggest that 1-hour incubation in modified Robbs buffer induces *wdb* RNAi oocytes to lose their arm cohesion and precociously enter anaphase 1. This result can be explained if PP2A protects cohesion on the chromosome arms.

To determine if cohesion of the centromeres was affected, we counted the number of centromere (CID or CENP-C) foci in the oocytes of each genotype. Each centromere detected by immunofluorescence during wild-type meiosis I contains two sister centromeres fused in a process that requires sister-chromatid cohesion (Wang et al., 2019). Thus, in wild-type oocytes we usually observed approximately eight centromere foci, as expected from four bi-oriented bivalents at metaphase I. In *wdb* RNAi oocytes, there was an increase in the number of oocytes with greater than 8 centromere foci, indicating that the sister chromatids were separating prematurely. In fact, 45% of *wdb* RNAi (*HMS01864* and *A5*) oocytes had >8 centromere foci, compared to 14% in wild-type (Figure 4D). This suggests that the sister centromeres are precociously separating, possibly due to loss of cohesion.

The sister centromere separation and precious anaphase phenotypes of *wdb* RNAi oocytes were relatively mild compared to other cohesion-defective mutants (Gyuricza et al., 2016; Wang et al., 2019). However, the severity of the spindle phenotypes was dramatically increased in both *wrd*^*GL00671*^; *wdb*^*HMS01864*^ and *wrd*^*-*^ ; *wdb*^*HMS01864*^ oocytes. The number of centromere foci was increased in *wrd*^*GL00671*^; *wdb*^*HMS01864*^ and *wrd*^*-*^ ; *wdb*^*HMS01864*^ oocytes (Figure 5B). Similarly, the precocious anaphase phenotype of chromosomes moving towards the poles was observed without a one-hour incubation in Robbs buffer (Figure 5C). These results can be explained if PP2A is required for the protection of sister chromatid cohesion during meiosis I.

**Figure 5:**
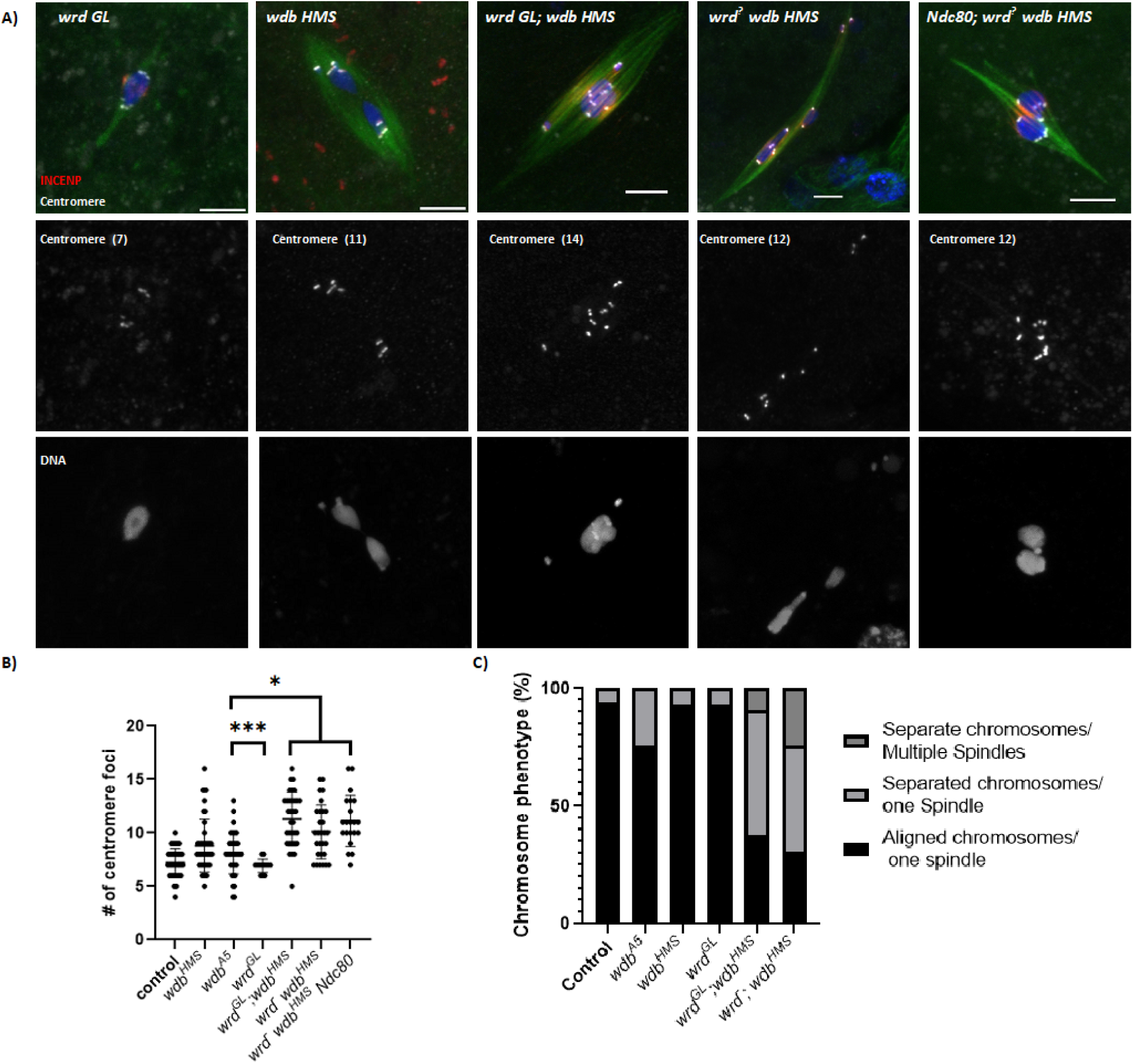
Precious anaphase in PP2A RNAi oocytes. (A) Stage 14 oocytes from single (*wrd*^*GL00671*^and *wdb*^*HMS01864*^) and double knockdown (*wrd*^*GL00671*^; *wdb*^*HMS01864*^ and *wrd*^*Δ*^ *wdb*^*HMS01864*^) females, with INCENP in red, centromeres in white, tubulin in green and DNA in blue. Centromeres and DNA are shown below the merged images and the number of centromere foci is indicated. Scale bars are 5μm. (B) Number of centromere foci in *wrd*^*GL00671*^ (n=13), *wdb*^*HMS01864*^ (n=43), *wrd*^*GL00671*^; *wdb*^*HMS01864*^ (n=33), *wrd*^*Δ*^ *wdb*^*HMS01864*^ (n=28) and *Ndc80; wrd*^*Δ*^ *wdb*^*HMS01864*^ (n=20) oocytes. (C) Percent chromosome phenotype for each genotype.

We previously found that two mechanisms can cause precocious sister centromere separation: loss of cohesion or inappropriate stabilization of kinetochore-microtubule (KT-MT) attachments (Wang et al., 2019). The inappropriate stabilization of KT-MT attachments depends on NDC80 and end on KT-MT attachments. To test the role of KT-MT attachments in the *wrd*^*-*^ ; *wdb*^*HMS01864*^ phenotype, we examined *wrd*^*-*^ ; *wdb*^*HMS01864*^; *Ndc80* RNAi oocytes. In *Ndc80* RNAi oocytes, end-on microtubule attachments are absent. The *wrd*^*-*^ ; *wdb*^*HMS01864*^; *Ndc80* RNAi oocytes exhibited both precocious anaphase and centromere separation, suggesting stabilization of end-on KT-MT attachments was not required for the *wrd*^*-*^ ; *wdb*^*HMS01864*^ phenotype. Because end-on attachments are necessary for separating sister centromeres in the presence of intact cohesins (Wang et al., 2019), observing centromere and homolog separation in the absence of end-on attachments is consistent with loss of cohesion in *wrd*^*-*^ ; *wdb*^*HMS01864*^ oocytes.

In addition to the precocious loss of cohesion, *wrd*^*GL00671*^; *wdb*^*HMS01864*^ or *wrd*^*-*^ ; *wdb*^*HMS01864*^ oocytes failed to form end on KT-MT attachments. In wild-type oocytes, most of the centromeres in stage 14 oocytes have end-on attachments, defined as when bundles of microtubules end at the centromeres (Figure 1A,C; Figure 2E; Figure 4A,C; Figure S 3). In contrast, *wrd*^*GL00671*^; *wdb*^*HMS01864*^ and *wrd*^*-*^ ; *wdb*^*HMS01864*^ oocytes had a high frequency of lateral microtubule attachments, defined when the centromeres are positioned along the side of microtubule bundles (Figure 5A, Figure S 3). Essentially all *wrd*^*GL00671*^; *wdb*^*HMS01864*^ and *wrd*^*-*^ ; *wdb*^*HMS01864*^ oocytes had this phenotype, which was rarely observed in wild-type oocytes. Assuming that lateral KT-MT attachments form first (Itoh et al., 2018; Shrestha et al., 2017), these data show that PP2A is required for the conversion from lateral to end-on attachments in oocytes. This conclusion helps explain why *Ndc80* RNAi had little effect on the *wrd*^*-*^ ; *wdb*^*HMS01864*^ oocyte phenotype. PP2A-B56 is required for stabilizing the end-on attachments that depend on NDC80. Because these PP2A-B56 depleted oocytes enter precocious anaphase, these results also suggest that lateral attachments are sufficient to move the chromosomes towards the poles, as also shown in *Drosophila* cell lines (Feijão et al., 2013).

### PP2A is required for bi-orientation of homologous chromosomes

To examine the effects of *PP2A* depletion on chromosome alignment and bi-orientation, we used fluorescent in situ hybridization (FISH). FISH probes were used that detected the pericentromeric regions of all three major chromosomes: the AACAC repeat (2nd chromosome), the Dodeca repeat (third chromosome) and the 359 repeat (X chromosome). In wild-type oocytes, correct bi-orientation is indicated when each pair of homologous centromeres separate towards opposite poles (Figure 6A). A low frequency of bi-orientation defects was observed in *tws* and *mts* RNAi oocytes (Figure 6B-C, G), showing that the TWS/ B55 subunit is not required for making the correct microtubule attachments during prometaphase I.

**Figure 6:**
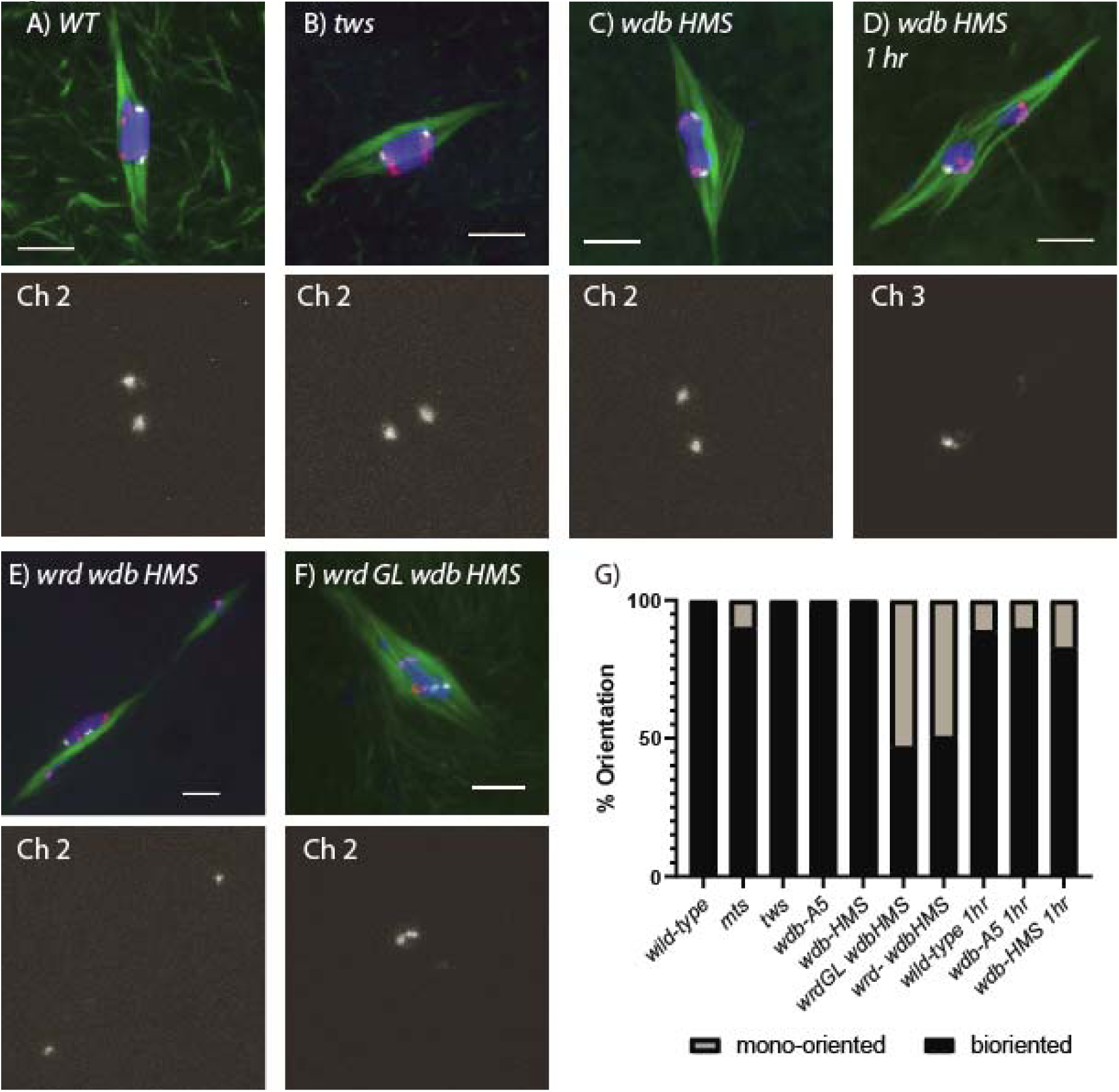
Analysis of bi-orientation in PP2A RNAi oocytes. FISH was performed with probes for the three major chromosomes in wild-type (WT) (A) and RNAi oocytes of (B) *tws*, (C) *wdb*^*HMS01864*^, (D) *wdb*^*HMS01864*^ incubated for 1 hour, (E) *wrd*^*Δ*^ *wdb*^*HMS01864*^, and (F) *wrd*^*GL00671*^ *wdb*^*HMS01864*^. The paracentric FISH probes were for the X chromosome (359bp repeat, Alexa594, purple), the 2nd chromosome (HET, Cy3, red) and the 3^rd^ chromosome (Dodeca, Cy5, white). An example of one FISH probe is shown in the lower panels (chromosome 2 or 3). Examples of mono-orientation are shown in D and F. Tubulin is in green, DNA in blue and the scale bars are 5μm. (G) Relative frequency of mono-oriented and bi-oriented centromeres in wild-type (n=117) and RNAi oocytes of *mts* (n=60), *tws* (n=72), *wdb*^*A5*^ (n=61), *wdb*^*HMS01864*^ (n=66), *wrd*^*GL00671*^ *wdb*^*HMS01864*^ (n=51), *wrd*^*Δ*^ *wdb*^*HMS01864*^ (n=73), wild-type incubated for 1 hour (n=45), *wdb*^*A5*^ incubated for 1 hour (n=98) and *wdb*^*HMS01864*^ incubated for 1 hour (n=87).

The frequency of bi-orientation defects in the *wdb*^*A5*^ and *wdb*^*HMS01864*^ RNAi oocytes was also low, which could have been due to redundancy with *wrd*. Indeed, oocytes depleted for both *wrd* and *wdb* had a much higher frequency of bi-orientation defects (Figure 6E-G). The FISH experiments also showed when cohesion had been lost on the chromosome arms. In wild-type metaphase I, pairs of homologous centromeres are separated but remain connected by chiasma and therefore within the same chromatin mass. In oocytes depleted for both *wrd* and *wdb*, pair of homologous centromeres were in different chromatin masses and had separated towards opposite poles (Figure 6E), indicating precocious anaphase due to loss of cohesion on the chromosome arms. Similarly, in *wdb* RNAi oocytes incubated for one hour in Robbs buffer, the FISH analysis showed that the two probe signals for each homolog were usually in different chromatin masses, showing that arm cohesion had been released, allowing the chromosomes to move towards the poles. In contrast to the oocytes depleted for both *wrd* and *wdb*, precocious anaphase was not associated with a high frequency of bi-orientation defects (Figure 6D).

### Spc105R Is Required for PP2A Localization

Because the function of kinases and phosphatases often depends on their localization, we examined the localization of WDB using either antibodies (Pinto and Orr-Weaver, 2017; Sathyanarayanan et al., 2004) or an HA-tagged transgene (Hannus et al., 2002). In wild-type oocytes, we found that WDB localizes prominently to the centromere regions (Figure 7A). In many oocytes, it was also possible to observe weaker localization to the chromosome arms (Figure 7B, Figure S 4B). Surprisingly, WDB protein was still detected at the centromeres using either the antibody or the HA-tagged transgene in *wdb* RNAi oocytes (Figure 7C, Figure S 4C). Thus, shRNA expression was effective enough to yield a mutant phenotype but did not eliminate WDB expression. One possibility is that the kinetochore localized WDB we observe is not sufficient for most PP2A-B56 functions.

**Figure 7:**
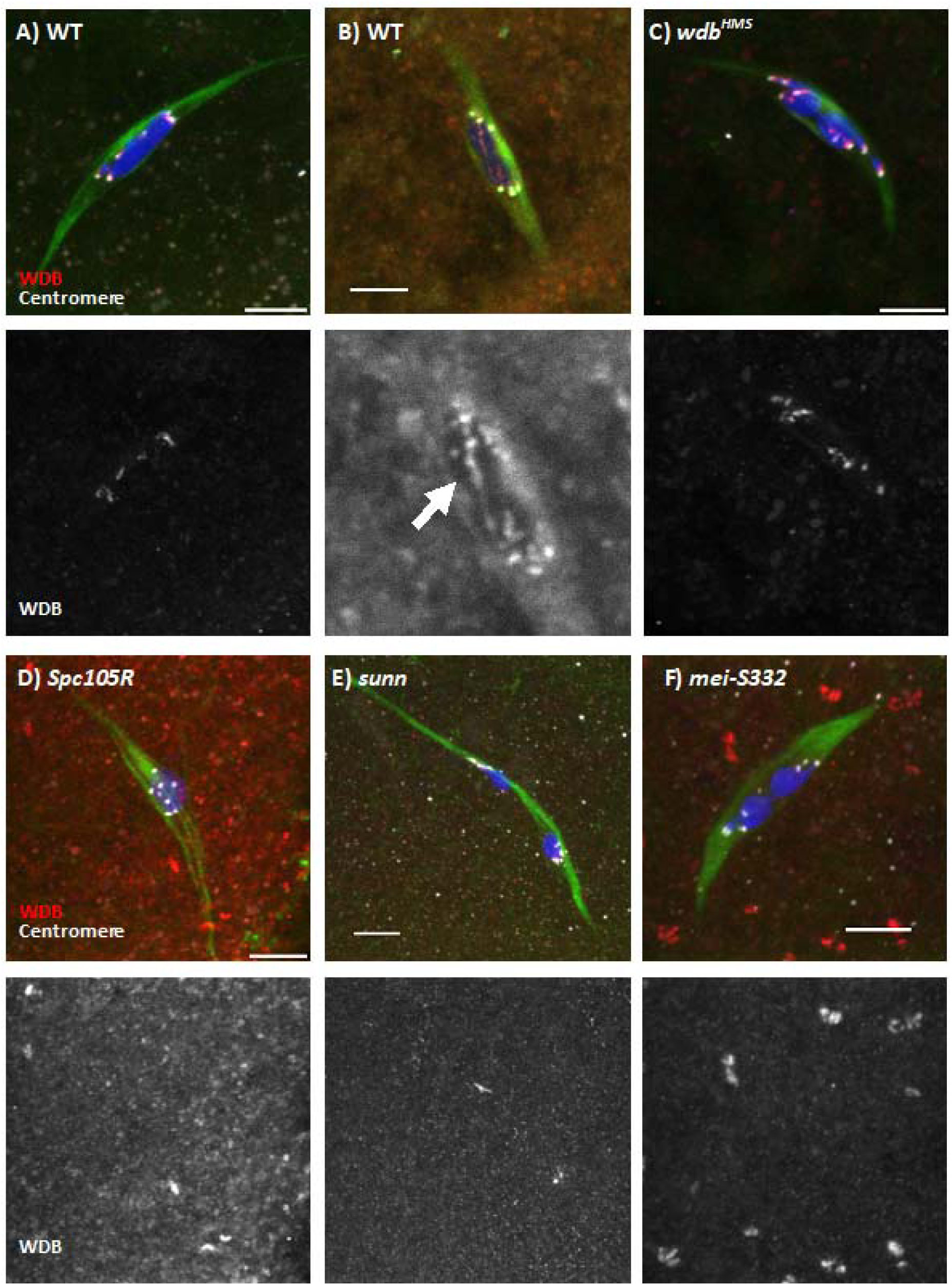
Localization of WDB in metaphase I of meiosis. Stage 14 oocytes showing WDB (red), centromeres (white), tubulin (green) and DNA (blue). The single channel images show WDB. Images show wild-type (WT) (A,B), RNAi (C,D) or homozygous mutant (E,F) oocytes. WDB was detected using a polyclonal antibody (Pinto and Orr-Weaver, 2017; Sathyanarayanan et al., 2004). The WDB channel in panel B is a higher magnification image to show threads of WDB signal (arrow), corresponding to PP2A on the chromosome arms. Scale bars are 5μm.

The strongest WDB accumulation colocalizes with centromere markers like CID/CENP-A or CENP-C, suggesting it is enriched in the centromere/ kinetochore regions. Therefore, we tested if WDB localization depends on SPC105R, which is a kinetochore protein previously shown to be required for sister centromere cohesion in meiosis I (Radford et al., 2015; Wang et al., 2019). *Spc105R* RNAi oocytes lacked WDB localization (Figure 7D), showing that Spc105R is required for the recruitment of WDB. To determine if cohesion is required for WDB localization, we used *sunn* mutant oocytes. SUNN is a stromalin-related protein required for sister chromatid cohesion in meiosis (Krishnan et al., 2014). WDB was present in at the centromeres in *sunn* mutant oocytes (Figure 7E), showing that cohesion is not required for WDB localization.

### Evidence for redundant mechanisms to recruit PP2A

In human cell lines, PP2A is recruited to the centromere regions by BUBR1 (Kruse et al., 2013; Xu et al., 2013) and Shugoshin (Kitajima et al., 2006; Riedel et al., 2006; Tang et al., 2006). Consistent with these studies, WDB was absent from the centromeres in *BubR1* RNAi oocytes (Figure S 4F). This was a surprising result, however, because *BubR1* RNAi oocytes do not have a cohesion defect, nor do they have a precocious anaphase phenotype (Wang et al., 2019). Similarly, WDB localization was reduced in *mei-S332* mutant oocytes, and also reduced in *mei-S332/+* heterozygotes (Figure 7F, Figure S 4D,E). *Drosophila mei-S332* mutants are viable, suggesting MEI-S332 is not required for cohesion in mitosis. Furthermore, chromosome segregation errors in *mei-S332* mutants primarily involve sister chromatids during meiosis II (Kerrebrock et al., 1995; Tang et al., 1998). Indeed, while MEI-S332 localizes to the centromere regions during meiosis I, the sister centromeres remain fused in *mei-S332* mutant oocytes (see below) (Kerrebrock et al., 1995; Moore et al., 1998). Therefore, while both BUBR1 and MEI-S332 are required to recruit WDB to the meiotic centromeres, unlike oocytes lacking PP2A, the absence of BUBR1 or MEI-S332 does not cause a meiosis I cohesion defect. In addition, *BubR1* RNAi oocytes were sensitive to BN2, suggesting that WDB does not have to ben enriched at that centromeres to destabilize the meiotic spindle (Figure S 5, see Discussion).

### Dalmatian May Protect Cohesion During Meiosis I

The absence of a strong meiosis I phenotype in *mei-S332* mutants could be explained if another protein recruits PP2A-B56 to the centromeres. Because we have only examined WDB localization, it is also possible this hypothetical pathway recruits WRD and not WDB. However, a *mei-S332; wrd* double mutant is viable (Pinto and Orr-Weaver, 2017), suggesting that in mitotic cells, WDB functions in the absence of MEI-S332. A candidate for a second protein that recruits PP2A is Dalmatian (DMT), which is a Soronin orthologue that has been proposed to recruit PP2A in *Drosophila* mitotic cells (Yamada et al., 2017). Consistent with this hypothesis, DMT colocalizes with MEI-S332 at the centromeres in metaphase I oocytes (Figure 8A). Mutations of *dmt* cause lethality (Salzberg et al., 1994). Therefore, to test the function of DMT in meiosis, we created shRNA lines to target *dmt* for tissue-specific RNAi. The strongest of the two shRNA lines caused sterility when expressed in oocytes with either *nosGal4* or *matαGal4* and lethality when expressed ubiquitously (Table 1, Table 2), suggesting the RNAi was effective. The strongest of these shRNAs reduced mRNA levels to 5% of wild-type levels. However, in these *dmt* RNAi oocytes, there was no centromere separation defects to indicate a loss of cohesion (Figure 8B,C,F). One explanation for the absence of a defect in *dmt* RNAi oocytes could be redundancy with MEI-S332. Therefore, we constructed *mei-S332* mutant females expressing *dmt* RNAi using *matαGal4*. These females, however, also had no centromere separation defects during meiosis I (Figure 8D-F).

**Figure 8:**
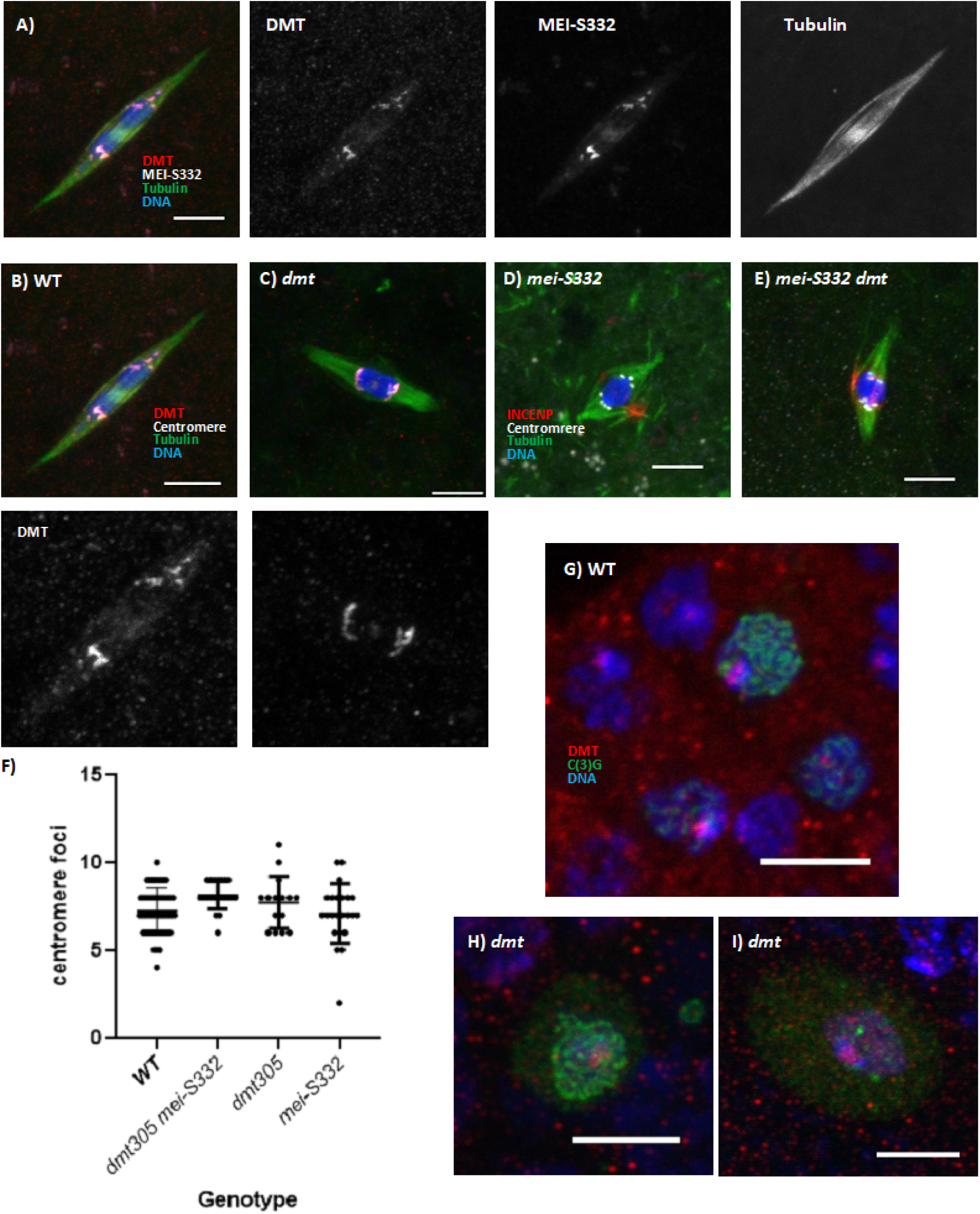
Localization of cohesion protection proteins in meiosis. (A) Wild-type (WT) oocyte showing DMT in red, MEI-S332 in white, tubulin in green and DNA in blue. Also shown are greyscale images of the DMT, MEI-SS332 and tubulin channels. Scale bar is are 5μm. (B,C) Wild-type and *dmt* RNAi oocytes showing with DMT in red, centromeres in white, tubulin in green and DNA in blue. DMT is shown in the bottom panel and scale bar is 5μm. (D-E) Phenotype of *dmt* RNAi and *mei-S332* mutant oocytes, with INCENP (red), centromeres (white), tubulin (green) and DNA (blue). (F) Summary of centromere foci in *dmt* RNAi (n=15), *mei-S332* mutant (n=23) or *dmt mei-S332* (n=21) oocytes. There are no significant differences between the three data sets by one-way ANOVA tests. However, *dmt mei-S332* is significantly different than *mei-S332* by a t-test (P=0.013). (G) DMT localization (red) in early prophase/pachytene oocytes. Oocytes are marked by C(3)G (green) along with DNA (blue). (H,I) DMT localization when the shRNA is expressed using *nosGAL4*. DMT localization is visible in these late stage pachytene (H) and prophase oocytes (I).

Another explanation for the absence of a meiotic defect in *dmt* RNAi oocytes is that the DMT protein is stable and loaded prior to expression of the shRNA by *matαGal4*, such as early meiotic S-phase or prophase. Consistent with this hypothesis, DMT localization to the centromeres was observed in *dmt* RNAi/ *matαGal4* oocytes (Figure 8C). Furthermore, we found that DMT localized to the meiotic chromosomes in the germarium, which contains oocytes early in prophase undergoing pachytene (Figure 8G-I). In contrast, we did not observe localization of WDB at these stages (Figure S 6). This result shows that DMT is loaded on the chromosomes prior to the domain of shRNA expression by *matαGal4*. Either DMT is stable enough to persist on the chromosomes without replenishment until metaphase I, or it is only loaded at a specific time in meiosis, such as pre-meiotic S-phase.

If DMT protein is loaded prior to the domain of *matαGal4* expression and stably maintained, shRNA expressed during S-phase and early pachytene by *nosGal4* should have a defect in meiosis. To test this idea, we examined RNAi of cohesin components, which are an example of proteins stably loaded onto the meiotic chromosomes in pre-meiotic S-phase (Gyuricza et al., 2016). Nondisjunction was observed when *nosGal4* was used to induce expression of shRNAs for cohesins *ord* (32%, n=1270) and *sunn* (15.6%, n=486). In contrast, nondisjunction was not observed with *matαGal4* and the same shRNAs for *ord* (0%, n=1622) or *sunn* (0%, n=547). These results can be explained if cohesins are stably maintained without new transcription throughout most of prophase. In contrast, expressing an shRNA to *mei-S332* with either *nosGal4* (40%, n=331) or *matαGal4* (27.8%, n=208) caused high levels of nondisjunction. These results show that *mei-S332* transcription is required late in prophase or during pro-metaphase I. If DMT behave like the cohesins, *nosGal4* should be necessary for *dmt* RNAi to affect meiosis. However, a loss of centromere cohesion was not observed in *dmt /nosGal4* RNAi oocytes (avg =7.9, n= 10). Finally, to test the hypothesis that MEI-S332 and DMT both recruit PP2A during meiosis I, we expressed *dmt* RNAi using *nosGal4* in *mei-S332* mutant females. These females were sterile, but unlike *dmt* RNAi /*nosGal4* females, very few oocytes were produced, which can be explained if DMT and MEI-S332 are redundant in the mitotic germline divisions. In summary, these data are consistent with the conclusion that *Drosophila* oocytes load cohesins and a protector of cohesin, DMT, during pre-meiotic S-phase.

## Discussion

Oocyte spindle assembly depends on the CPC. Inhibiting Aurora B in oocytes with either Binucleine 2 (this paper) or RNAi (Radford et al., 2012b) results in total loss of the meiotic spindle. This study was initiated to identify and characterize the phosphatases that antagonize Aurora B. In the process of this analysis, we investigated the role of PP2A in oocyte meiosis.

### Initiation and maintenance of the meiotic spindle

Knockdown of PP2A components suppressed the BN2-induced loss of the meiotic spindle. The implication is that constant Aurora B activity is required to maintain the spindle, and in its absence, PP2A removes activating phosphorylation events on meiotic spindle proteins. This was shown directly using phosphorylated INCENP, which is a target of Aurora B. Based on indirect evidence, kinetochore protein SPC105R and kinesins KLP10A and Subito also depend on this balance of phosphorylation. Surprisingly, depletions of both PP2A subtypes, B55 (e.g. TWS) and B56 (e.g. WDB), suppressed BN2-induced spindle loss. These two complexes are usually targeted to different substrates. These results can be explained if PP2A-B55 depletion reduces the activity of PP2A-B56. This is plausible because B56 activity may depend on Polo kinase, which in turn may be regulated by B55 (Rangone et al., 2011; Wang et al., 2011b). Alternatively, PP2A-B55 may directly regulate Aurora B activity via CDK1 (Hummer and Mayer, 2009; Kitagawa et al., 2014).

In CPC depleted *Xenopus* egg extracts, microtubule stabilization around chromatin fails, whereas when both the CPC and Kinein-13 MCAK are depleted, microtubules are stabilized (Sampath et al., 2004). Similarly, we found that depletion of Kinesin-13 KLP10A partially alleviated the spindle loss in BN2 treated oocytes. These results indicate that the CPC negatively regulates spindle depolymerizing factors. In contrast, the absence of spindle assembly in CPC RNAi oocytes is not suppressed by simultaneous knockdown of Kinesin-13 KLP10A (Radford et al., 2012a; Radford et al., 2012b). The different effects of KLP10A knockdown on *aurB* RNAi and BN2 inhibition is most likely due to the timing of CPC depletion. In RNAi oocytes, loss of a Kinesin-13 does not restore spindle assembly because factors required for the initiation of spindle assembly are not activated. In contrast, spindle disassembly in oocyte treated with BN2 depends on depolymerizing activities, and therefore requires Kinesin-13.

PP2A-B56 probably regulates factors which maintain spindle integrity. Subito localization depends on the CPC and, along with the kinesin-11 NCD, may depend on high levels of Aurora B near the chromosomes to be activated (Beaven et al., 2017; Das et al., 2018). Interestingly, Subito contains one predicted PP2A-B56 binding motif (FDNIQESEE) (Hertz et al., 2016), and it is in a region we proposed negatively regulates Subito activity (Das et al., 2018). In addition to these two kinesins, it is likely that several spindle assembly factors are regulated by antagonism between Aurora B and PP2A. This is probably a conserved activity, as PP2A has been shown to oppose Aurora B activity in the central spindle of HeLa cells (Bastos et al., 2014).

### PP2A-B56 is required to maintain meiotic cohesion

In addition to the role of antagonizing Aurora B in spindle maintenance, PP2A-B56 is required for two additional meiosis I processes, sister chromatid cohesion and stabilization of end-on KT-MT attachments. The strongest sister chromatid cohesion phenotypes were observed when both B56 subunits, WDB and WRD, were simultaneously knocked out. The loss of cohesion resulted in the separation of sister centromeres and, as expected from the loss of arm cohesion (McKim et al., 1993) precocious anaphase. PP2A-B56 maintains cohesion by dephosphorylation of cohesin subunits, preventing Separase from cleaving the Kleisin subunit (Gutierrez-Caballero et al., 2012). In many cell types, PP2A-B56 is recruited by Shugoshin. Indeed, we observed a reduction in WDB localization in a *mei-S332* (the *Drosophila* Shugoshin homolog) mutant. However, there is a striking difference between the phenotype from loss of PP2A-B56 and MEI-S332, even though the latter is required for its localization. The lack of a meiosis I cohesion defect in *mei-S332* mutants can be explained if a low level of PP2A-B56 is recruited in its absence. Because these are null alleles of *mei-S332*, this low level of PP2A-B56 must depend on a MEI-S332-independent mechanism for recruitment to the chromosomes. Similar evidence for a second pathway to recruit PP2A was found in *Drosophila* male meiosis (Pinto and Orr-Weaver, 2017). While WDB localization was observed in both prophase I and metaphase I spermatocytes, it was eliminated only from the metaphase I spermatocytes in *mei-S332* mutants. Thus, the prophase I localization of WDB did not depend on MEI-S332.

The *Drosophila* Soronin homolog Dalmatian (DMT), and not MEI-S332, is required for cohesion in *Drosophila* cells (Yamada et al., 2017) and has sequence features, including the LxxIxE motif, that suggest it recruits PP2A-B56 (Hertz and Nilsson, 2017). However, a loss of cohesion phenotype was not observed in *dmt* RNAi oocytes. There are two possible explanations for these results. First, DMT, like the other cohesins required for cohesion, is a stable protein that is only loaded onto chromosomes during pre-meiotic S-phase (Gyuricza et al., 2016). Indeed, we observed DMT localization early in meiotic prophase. Second, DMT and MEI-S332 may redundantly protect cohesion in oocytes. We propose that PP2A-B56 recruitment by DMT occurs during S-phase and ensures protection during the long oocyte prophase and into metaphase I. MEI-S332/SGO recruitment of PP2A-B56 is established late, once the nuclear envelope breaks down, and also protects cohesion during the meiotic divisions. Testing this model requires a precise knock out DMT, during premeiotic S-phase but after the mitotic divisions of the germline.

### PP2A-B56 regulates kinetochore-microtubule attachments

When both B56 subunits, WDB and WRD, were knocked out, we observed loss of end-on KT-MT attachments and bi-orientation defects. The bi-orientation errors could be a consequence of the attachment defects. PP2A, possibly via recruitment by BUBR1, can stabilize KT-MT attachments in mitotic cells (Foley et al., 2011; Kruse et al., 2013; Suijkerbuijk et al., 2012; Xu et al., 2013) and meiotic cells (Tang et al., 2016). Similarly, in *Drosophila* oocytes, we have shown that BubR1 stabilizes KT-MT attachments (Wang et al., 2019). In the absence of end-on attachments, lateral attachments persist (Feijão et al., 2013; Radford et al., 2015). The target of PP2A in this process could be the N-terminal domain of SPC105R (Nijenhuis et al., 2014; Smith et al., 2019). The N-terminal domain of SPC105R/KNL1 has two known properties, it binds microtubules (Bajaj et al., 2018; Espeut et al., 2012) and it a contains two Aurora B phosphorylation sites (Liu et al., 2010; Rosenberg et al., 2011; Welburn et al., 2010). Furthermore, Aurora B activity can inhibit the conversion of lateral to end-on KT-MT attachments (Kalantzaki et al., 2015; Shrestha et al., 2017). Based on these observations, we propose that Aurora B kinase phosphorylation of SPC105R promotes lateral attachments, and PP2A reverses these events to allow end-on attachments to occur.

### Multiple independent pools of PP2A-B56 regulate meiosis

PP2A-B56 activity has two important functions at the centromeres or kinetochores: protecting cohesion and stabilizing KT-MT attachments. The dramatic disorganization of chromosomes and spindle, and precocious entry into anaphase, in PP2A-B56 depleted oocytes is a consequence of these two defects. PP2A localization depends on BubR1 and MEI-S332. However, the PP2A-B56 loss of function phenotypes are much stronger than BUBR1 or MEI-S332 loss of function. We propose there are at least three pools of PP2A-B56 in oocyte meiosis. The first pool is defined by localization of WDB that depends on BUBR1 and MEI-S332. Based on the mild phenotype of BUBR1 RNAi or *mei-S332* mutant oocytes, this pool is not essential for cohesion but could regulate end-on attachments. The second pool is defined as being required for cohesion, is independent of BUBR1, may include the weak chromosome arm localization observed in many of our oocyte images and could depend on Dalmatian. The two pools of PP2A that regulate cohesion and KT-MT attachments both depend on SPC105R. The third pool is defined by being required for antagonizing the spindle assembly function of Aurora B. This pool does not depend on kinetochore localization, as shown by the sensitivity of *Spc105R* (Wang et al., 2019) and *BubR1* RNAi (this paper) oocyte spindles to BN2 treatment.

An important question for future studies is how different pools of PP2A-B56 are independently regulated and interact. Prior studies have suggested that BubR1 (Kruse et al., 2013; Suijkerbuijk et al., 2012; Xu et al., 2013) and Shugoshin/ MEI-S332 (Kitajima et al., 2006; Riedel et al., 2006; Tang et al., 2006) regulate two distinct PP2A-B56 pools. In contrast, our data suggests BUBR1 and MEI-S332 recruit one pool while SPC105R recruits two functionally distinct pools of PP2A-B56. It will be interesting to determine how the function of these two kinetochore pools are separated and what recruits the second pool. Additionally, it remains to be determined if the non-kinetochore PP2A-B56 pool regulates kinetochore function, as proposed in mouse oocytes (Touati et al., 2015). And if so, it becomes critical to understand what recruits PP2A to the microtubules of the meiotic spindle. Understanding how PP2A-B56 achieves its multiple functions will require connecting specific functions with its spatial regulation.

## Materials and Methods

### Tissue specific knockdowns using expression of transgenes and shRNAs

The UAS/Gal4 system was used for tissue-specific expression of transgenes and shRNAs. In most experiments the transgenes and shRNAs, under UAS control, were expressed using *P{w*^*+mC*^*=matalpha4-GAL-VP16}V37* (*mataGAL4*), which induces expression after pre-meiotic DNA replication and early pachytene, and persists throughout most stages of meiotic prophase during oocyte development in *Drosophila* (Radford et al., 2012b; Sugimura and Lilly, 2006). For expression throughout the germline, including the germline mitotic divisions and early meiotic prophase, we used *P{w*^*+mC*^*=GAL4::VP16-nos*.*UTR}MVD1* (*nos-GAL4-VP16*). For ubiquitous expression, we used *P{tubP-GAL4}LL7*. The RNAi lines used in this study are listed in Table 1. We first selected RNAi lines that caused lethality when the shRNA was under the control of *P{tubP-GAL4}LL7* and sterile under the control of *mataGAL4*. Two RNAi lines were not used because of weak phenotypes. *HSM1804* (*mts*) was fertile with *mataGAL4* and *HMS1921* (*PP2A*-*29B*) was viable with *P{tubP-GAL4}LL7*.

The *wrd*^*Δ*^ mutant was generated by FLP-mediated site-specific recombination that removed most of the coding region (Moazzen et al., 2009). The deletion *Df(3R)189* was made by imprecise excision of a P-element within the *wrd* gene and removes all of *wrd* and a couple genes on each side (Viquez et al., 2006). For knockdown of *wrd* and *wdb*, mutations and shRNA were combined to generate *Df(3R)189 mataGAL4/ wrd*^*Δ*^ *wdb*^*HMS01864*^ females, or two shRNAs were combined to generate *GL00671*/+; *mataGAL4/ wdb*^*HMS01864*^ females.

### Generation of shRNA lines and analysis by RT-PCR

Sequences for shRNAs targeting *wdb* and *dmt* were designed using DSIR (http://biodev.extra.cea.fr/DSIR/DSIR.html) (Vert et al., 2006) and the GPP Web Portal (http://www.broadinstitute.org/rnai/public/seq/search). These were cloned into the pVallium22 vector for expression under control of the UASP promoter.

To measure the knockdown of mRNA in oocytes, Taqman qRT-PCR was used. In order to extract RNA from oocytes of interest, female flies were placed in yeasted vials for approximately 3 days. The females were then grinded in a blender containing 1x PBS, and oocytes were filtered through meshes, as described below for cytological analysis of stage 14 oocytes. 50 mg of oocytes were weighed out, and 1mL of TRIzol Reagant was added. RNA extraction was completed, according to manufacturer’s instructions (ThermoFisher Scientific). A nanodrop was used to measure the concentration of the RNA, and then 2 μg was used in to prepare cDNA using a High Capacity cDNA Reverse Transcription Kit (Applied Biosystems). After using a nanodrop to determine the concentration of cDNA in the sample, qPCR was performed using a TaqMan Assay (Applied Biosystems) and four replicates per reaction.

### Cytology of Stage 14 Oocytes and Drug treatment

To prepare oocytes for cytology, 0-3 day old females were placed in yeasted vials for 1-2 days as described (Gilliland et al., 2009). The females were grinded in a blender with modified Robbs buffer and filtered through a series of meshes to separate the stage 14 oocytes from other body parts. It was at this stage that oocytes were treated with 50 µM BN2 in modified Robbs for one hour. The BN2 was dissolved in DMSO to make a 50mM stock solution and then 1 ul was added to 999ul of modified Robbs. Control oocytes were incubated for 1 hr in 1ml of modified Robbs buffer containing 1ul DMSO.

Oocytes were fixed in fixation buffer with 5% formaldehyde and heptane and then washed with 1X PBS (Radford and McKim, 2016). Oocyte membranes were mechanically removed by rolling the oocytes between the frosted side of a glass slide and a coverslip. The rolled oocytes were then rinsed into 15mL conical tubes containing PBS/1% Triton X-100 and were rotated for 1.5 to 2 hours. Oocytes were washed in PBS/0.05% Triton X-100 and subsequently transferred to a 1.5 ml Eppendorf tube. Oocytes were then blocked in PTB for 1 hour and then incubated with primary antibodies overnight at 4 °C. The next day, the oocytes were washed 4 times in PTB and the secondary antibody was added. After incubating at room temperature for 4 hours, the oocytes were washed in PTB, stained for DNA with Hoechst33342 (10 µg/ml) and then washed 2X in PTB.

The primary antibodies for this study were tubulin monoclonal antibody DM1A (1:50), directly conjugated to FITC (Sigma, St. Louis). rat anti-Subito (Jang et al., 2005), rat anti-INCENP (Wu et al., 2008), rabbit anti-phospho-INCENP (Salimian et al., 2011; Wang et al., 2011a), guinea pig anti-MEI-S332 (Moore et al., 1998), rabbit anti-CENP-C (Heeger et al., 2005), rabbit anti-SPC105R (Schittenhelm et al., 2007), rabbit anti-WDB (Sathyanarayanan et al., 2004), rabbit anti-DMT (Yamada et al., 2017), rat anti-HA (Roche) and rabbit anti-CID (Active Motif), mouse anti-C(3)G (Page and Hawley, 2001). These primary antibodies were combined with either a Cy3, Alexa 543, Cy5 or Alexa 647 secondary antibody pre-absorbed against a range of mammalian serum proteins (Jackson Immunoresearch and ThermoFisher). This protocol was modified for FISH as described (Radford and McKim, 2016) using probes corresponding to the X chromosome 359 repeat labeled with Alexa 594, 2^nd^ chromosome AACAC repeat labeled with Cy3 and the 3^rd^ chromosome dodeca repeat labeled with Cy5 (IDT). The oocytes were mounted in Slowfade Gold antifade (ThermoFisher) and imaged using a Leica TCS SP8 confocal microscope with a 63X, N.A. 1.4 lens.

### Image analysis, quantification and statistical analysis

Leica Confocal Software was used to create maximum projections of complete image stacks of individual or merged channels. Image cropping and addition of scale bars was done in Adobe Photoshop. Centromere foci and spindle intensity were measured using Imaris image analysis software (Bitplane). Statistical tests were performed using GraphPad Prism software. All the numbers of the centromere foci or chromosome phenotypes were pooled together and ran one-way ANOVA followed by post hoc pairwise Tukey’s multiple comparison test. Details of statistical evaluations and the numbers of samples are provided in the figure legends.

### Nondisjunction/Fertility Assay

*Drosophila* crosses were used to determine the rate of nondisjunction and fertility of certain genotypes. In this cross, female virgin flies with a gene of interest were mated to *y w/B*^*s*^*Y* males. These males carry a dominant *Bar* mutation on the Y chromosome. Therefore, the progeny from a cross with *y w/B*^*s*^*Y* males are Bar males (XY) and wild-type females (XX). Nondisjunction of the sex chromosomes during meiosis results in four different zygotes, two are inviable (OY, XXX) and two are viable with distinguishing phenotypes, Bar females (XXY) and wild-type males (XO). The nondisjunction rate was calculated by (2*NDJ/(normal progeny + 2NDJ).

Figure S 1: BN2 treatment in *Pp1-87B* RNAi oocytes. (A) Wild-type (WT) and *Pp1-87B* RNAi oocytes treated with either 0.001% DMSO or 25µM BN2 immunostained with DNA (blue), tubulin (green), pINCENP (red) and INCENP (white). All images are maximum projections of Z-stack and scale bars are 5 µm. (B) Frequency of spindle loss in wild-type (n=41 and 35) and *Pp1-87B* RNAi (n=114 and 144) oocytes, treated with DMSO or BN2. Error bars show SEM of each category. ***=P<0.001 (Fisher’s exact test). (C) Kinetochore (SPC105R) localization (red) in *Pp1-87B* RNAi oocytes treated with either DMSO (n=6) or BN2 (n=8). In the merged image, SPC105R is shown in red along with DNA (blue) and tubulin (green). (D) Subito localization (red) in *Pp1-87B* RNAi oocytes treated with either DMSO (n=12) or BN2 ((n=8).

Figure S 2: FRAP analysis comparing the recovering time of wild-type and *mts* RNAi oocytes expressing *GFPS65C-alpha-Tub84B*.

Figure S 3: Stage 14 oocytes showing end-on and lateral microtubule attachments. For each image, a higher magnification image shows examples of end on attachments in wild-type (A) and lateral attachments (B-D) in *wrd wdb* double knockout oocytes. Oocytes are shown with DNA in blue, tubulin in green, centromeres in white and either WDB (A) or INCENP (B-D) in red. All images are maximum projections of Z-stack and scale bars are 5 µm.

Figure S 4: Additional images of WDB localization using an HA-tagged transgene (Hannus et al., 2002) showing DNA in blue, tubulin in green, HA-WDB in red, and centromeres in white. Arrow in panel B shows a thread of WDB between the chromosomes. All images are maximum projections of Z-stack and scale bars are 5 µm.

Figure S 5: PP2A antagonizes Aurora B activity in *BubR1* RNAi oocytes. Oocytes were treated with 50µM BN2 and are shown with INCENP in red, tubulin in green, centromeres in white and DNA in blue. Scale bars are 5μm. (A) A BN2 treated *wild-type* oocyte lacks spindle microtubules (n=9/10). (B-C) Three representative examples of BN2 treated *BubR1 RNAi* oocytes showing a severe reduction in spindle microtubules (n=10/10).

Figure S 6: WDB localization is not observed in meiotic prophase. WDB-HA (red) is not concentrated at the centromeres (white) in early prophase (A – the germarium) and mid-prophase (B,C – vitellarium oocytes). ORB (green) is a cytoplasmic protein that is enriched in the oocytes (Lantz et al., 1994). Scale bar is 5 µm.

