## Supplemental Figures for "Multiple pools of Protein Phosphatase 2A-B56 function to antagonize spindle assembly, promote kinetochore attachments and maintain cohesion in *Drosophila* Oocytes"

Figure S 1: BN2 treatment in *Pp1-87B* RNAi oocytes. (A) Wild-type (WT) and *Pp1-87B* RNAi oocytes treated with either 0.001% DMSO or 25µM BN2 immunostained with DNA (blue), tubulin (green), pINCENP (red) and INCENP (white). All images are maximum projections of Z-stack and scale bars are 5 µm. (B) Frequency of spindle loss in wild-type (n=41 and 35) and *Pp1-87B* RNAi (n=114 and 144) oocytes, treated with DMSO or BN2. Error bars show SEM of each category. ***=P<0.001 (Fisher's exact test). (C) Kinetochore (SPC105R) localization (red) in *Pp1-87B* RNAi oocytes treated with either DMSO (n=6) or BN2 (n=8). In the merged image, SPC105R is shown in red along with DNA (blue) and tubulin (green). (D) Subito localization (red) in *Pp1-87B* RNAi oocytes treated with either DMSO (n=12) or BN2 ((n=8).

Figure S1

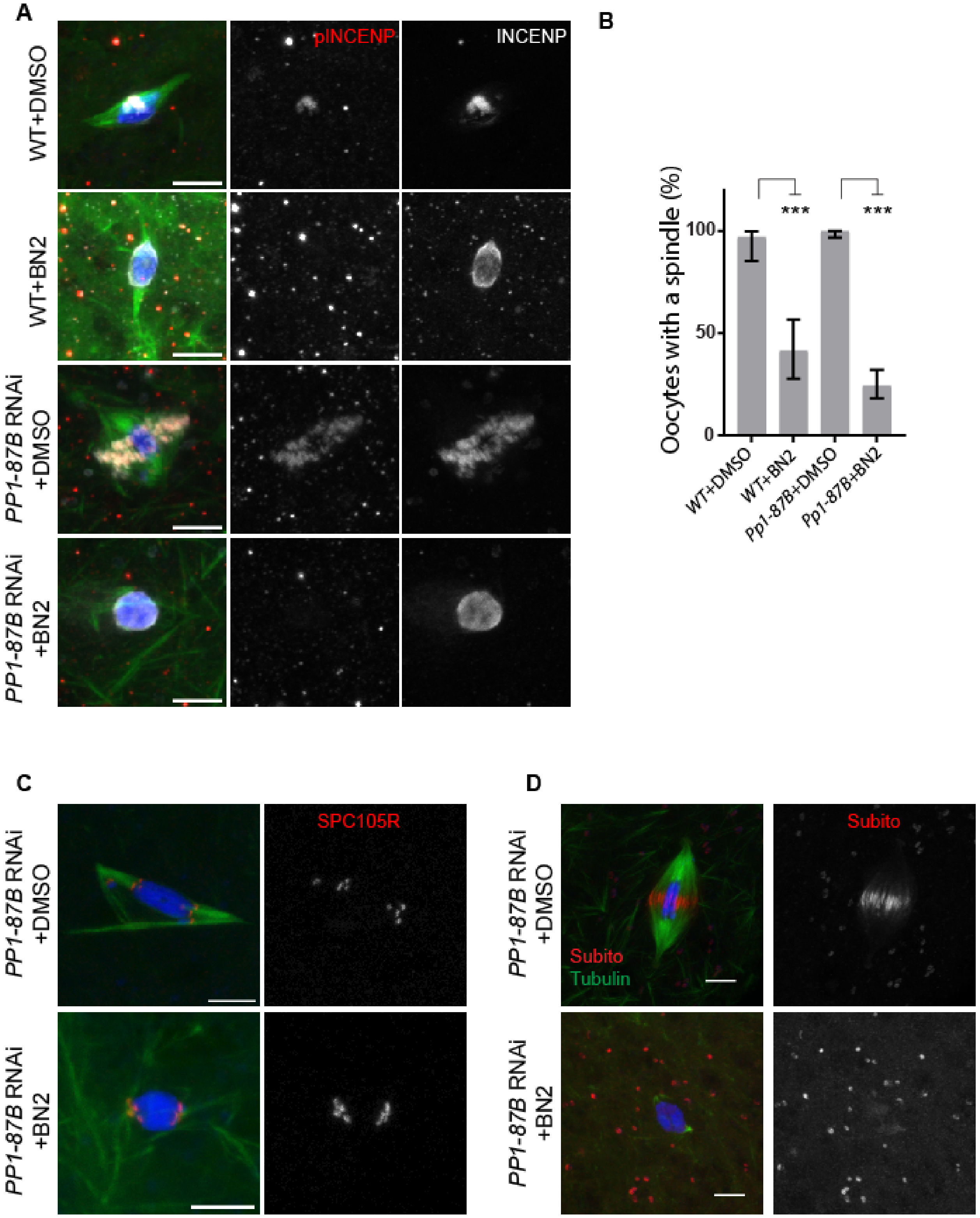
A
WT+DMSO
WT+BN2
PP1-87B RNAi +DMSO
PP1-87B RNAi +BN2
pINCENP
INCENP
B
Oocytes with a spindle (%)
100
50
0
***
***
WT+DMSO
WT+BN2
Pp1-87B+DMSO
Pp1-87B+BN2
C
PP1-87B RNAi +DMSO
PP1-87B RNAi +BN2
SPC105R
D
PP1-87B RNAi +DMSO
PP1-87B RNAi +BN2
Subito
Subito
Tubulin

Figure S 2: FRAP analysis comparing the recovering time of wild-type and *mts* RNAi oocytes expressing *GFPS65C-alpha-Tub84B*.

Figure S2

| | **Control** | ***mts*** |
|---|---|---|
| Spindle 1 | 4.6 s | 8.42 s |
| Spindle 2 | 9.94 s | 4.59 s |
| Spindle 3 | 16.45 s | 7.9 s |
| Spindle 4 | 8.87 s | 13.06 s |
| Spindle 5 | 8.21 s | 7.91 s |
| Spindle 6 | 8.41 s | |
| Average half-life | 9.41 s | 8.38 s |

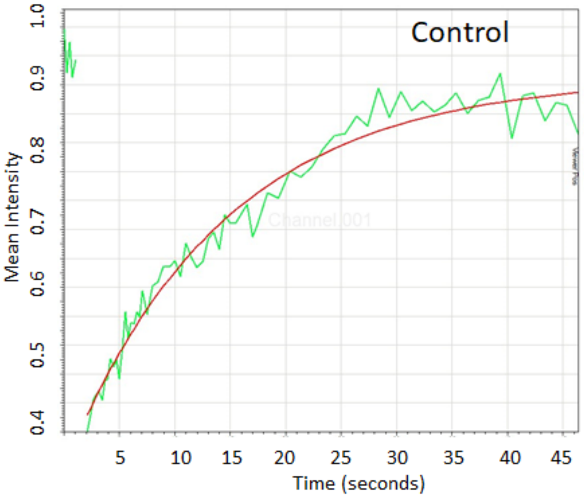

Figure S 3: Stage 14 oocytes showing end-on and lateral microtubule attachments. For each image, a higher magnification image shows examples of end on attachments in wild-type (A) and lateral attachments (B-D) in *wrd wdb* double knockout oocytes. Oocytes are shown with DNA in blue, tubulin in green, centromeres in white and either WDB (A) or INCENP (B-D) in red. All images are maximum projections of Z-stack and scale bars are 5 µm.

Figure S3

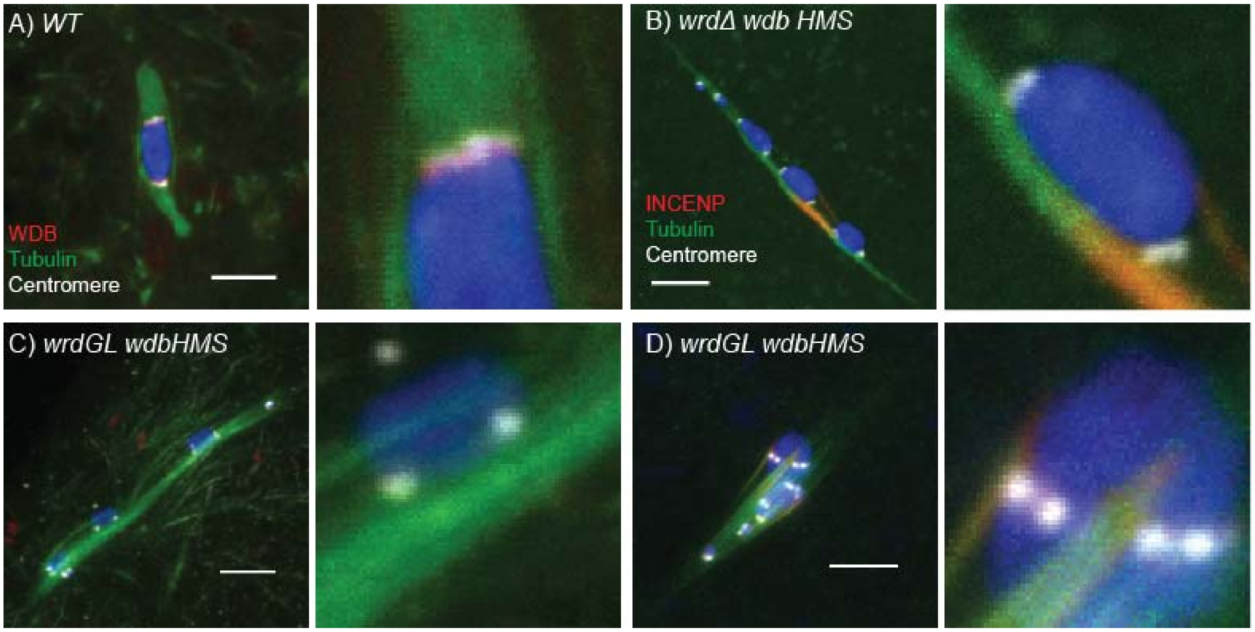

Figure S 4: Additional images of WDB localization using an HA-tagged transgene (Hannus et al., 2002) showing DNA in blue, tubulin in green, HA-WDB in red, and centromeres in white. Arrow in panel B shows a thread of WDB between the chromosomes. All images are maximum projections of Z-stack and scale bars are 5 μm.

Figure S4

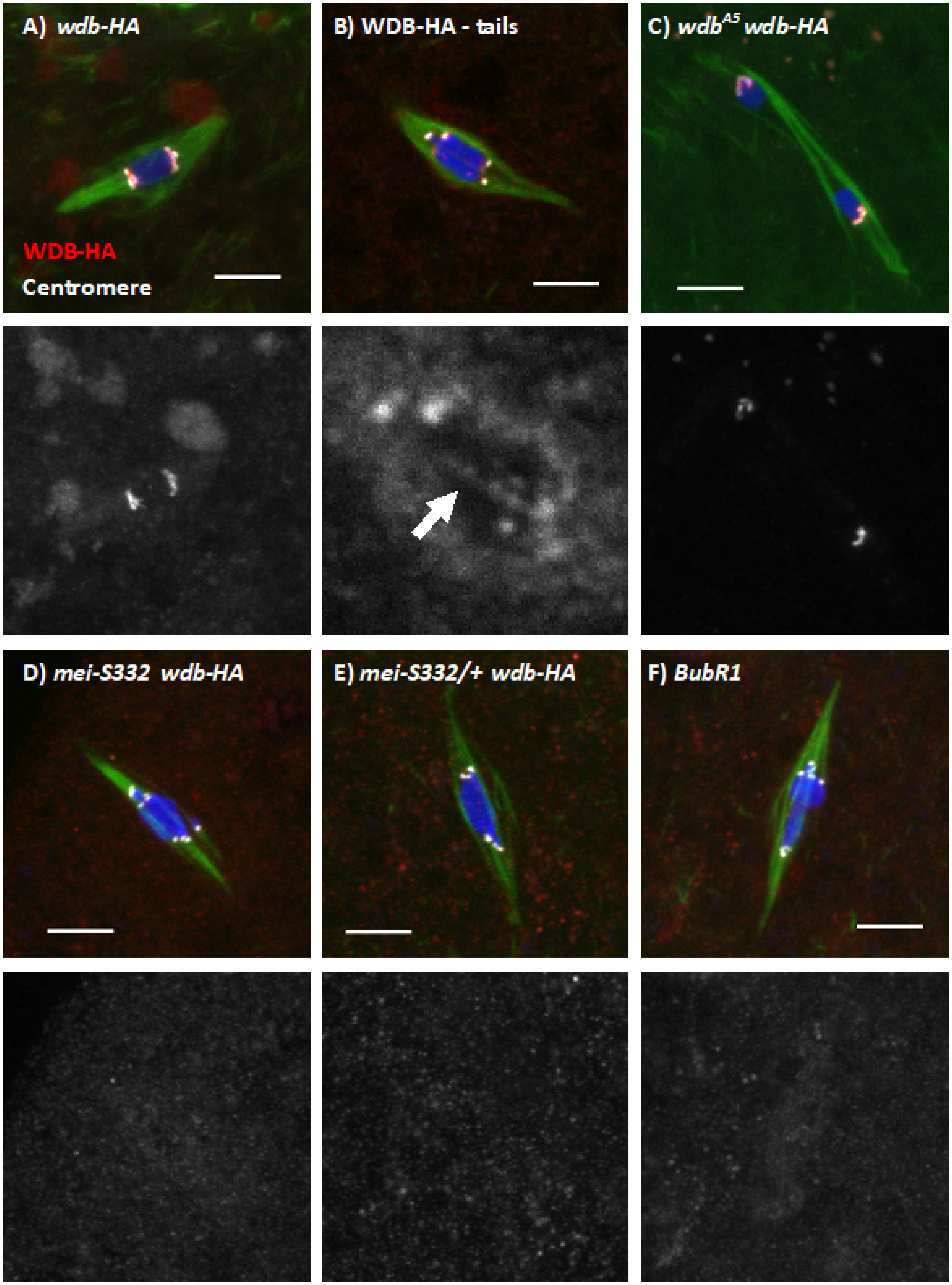
A) wdb-HA
WDB-HA
Centromere
B) WDB-HA - tails
C) wdb^A5 wdb-HA
D) mei-S332 wdb-HA
E) mei-S332/+ wdb-HA
F) BubR1

Figure S 5: PP2A antagonizes Aurora B activity in *BubR1* RNAi oocytes. Oocytes were treated with 50µM BN2 and are shown with INCENP in red, tubulin in green, centromeres in white and DNA in blue. Scale bars are 5µm. (A) A BN2 treated *wild-type* oocyte lacks spindle microtubules (n=9/10). (B-C) Three representative examples of BN2 treated *BubR1 RNAi* oocytes showing a severe reduction in spindle microtubules (n=10/10).

Figure S5

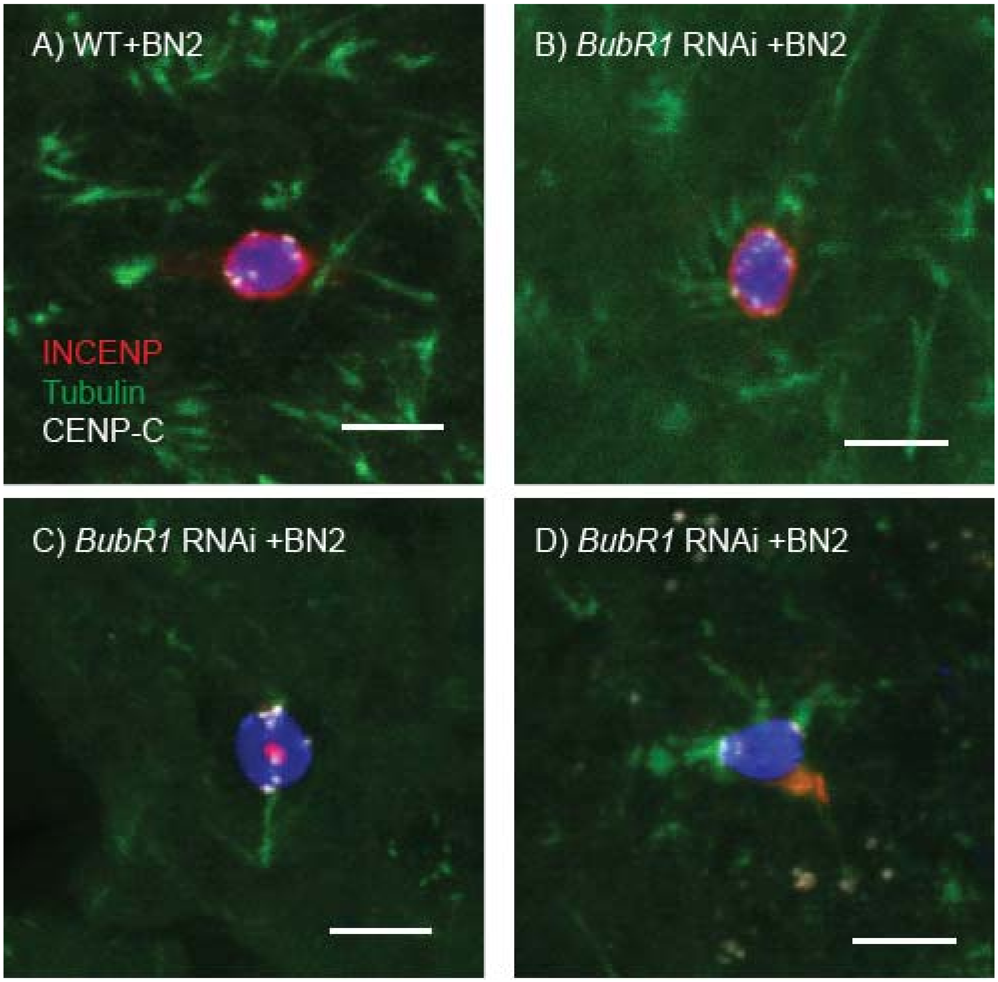

Figure S 6: WDB localization is not observed in meiotic prophase. WDB-HA (red) is not concentrated at the centromeres (white) in early prophase (A – the germarium) and mid-prophase (B,C – vitellarium oocytes). ORB (green) is a cytoplasmic protein that is enriched in the oocytes (Lantz et al., 1994). Scale bar is 5 µm.

Figure S6

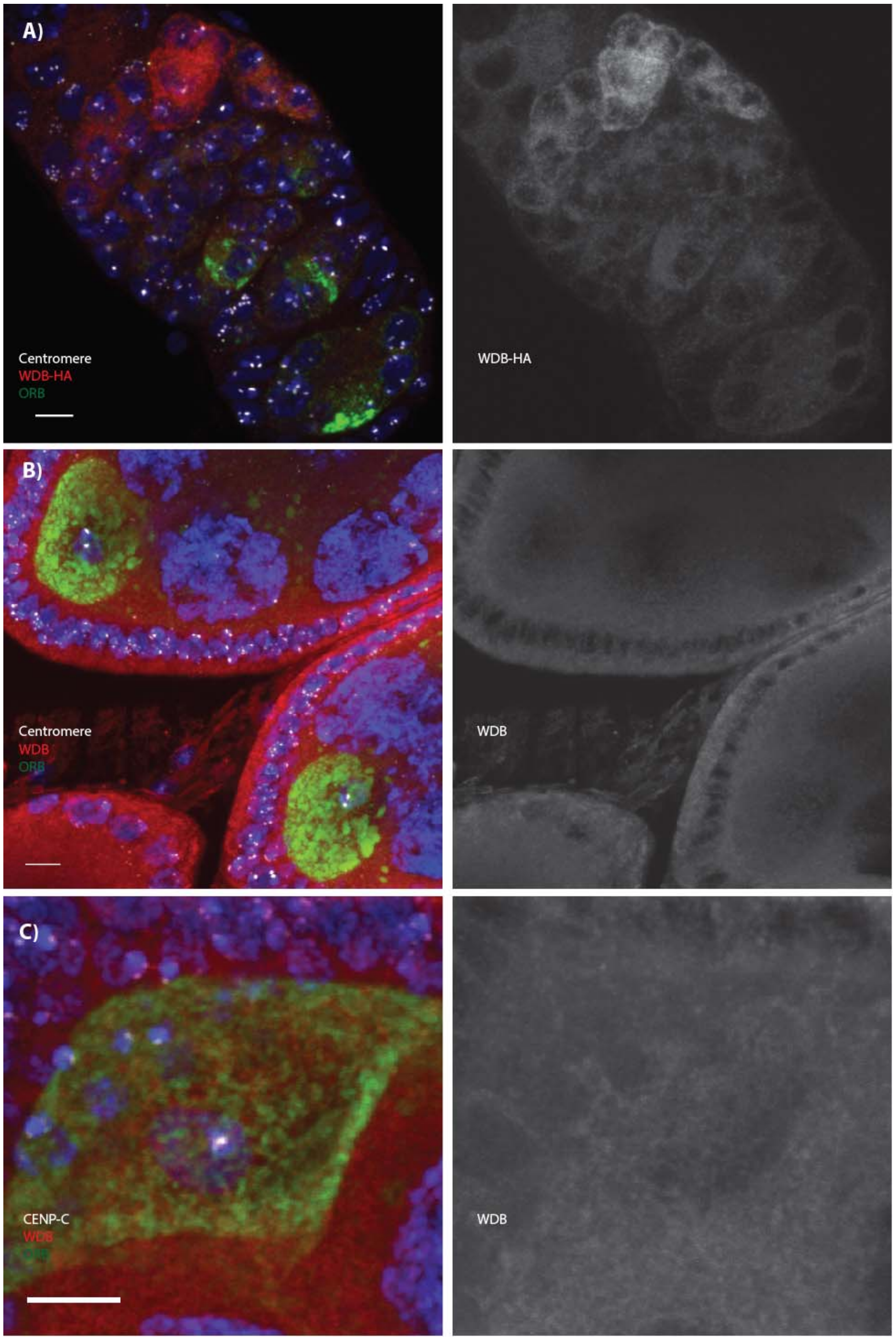
A)
Centromere
WDB-HA
ORB
WDB-HA
B)
Centromere
WDB
ORB
WDB
C)
CENP-C
WDB
WDB
